# Pivotal roles of *Plasmodium falciparum* lysophospholipid acyltransferase 1 in cell cycle progression and cytostome internalization

**DOI:** 10.1101/2024.01.13.575495

**Authors:** Junpei Fukumoto, Minako Yoshida, Suzumi M. Tokuoka, Eri Saki H. Hayakawa, Shinya Miyazaki, Takaya Sakura, Daniel Ken Inaoka, Kiyoshi Kita, Jiro Usukura, Hideo Shindou, Fuyuki Tokumasu

## Abstract

The rapid intraerythrocytic replication of *Plasmodium falciparum*, a deadly species of malaria parasite, requires a quick but constant supply of phospholipids to support marked cell membrane expansion. In the malarial parasite, many enzymes functioning in phospholipid synthesis pathway have not been identified or characterized. Here, we identified *P. falciparum* lysophospholipid acyltransferase 1 (PfLPLAT1) and showed that PfLPLAT1 is vital for asexual parasite cell cycle progression and cytostome internalization. Deficiency in PfLPLAT1 resulted in decreased parasitemia and prevented transition to the schizont stage. Parasites lacking PfLPLAT1 also exhibited distinctive omega-shaped vacuoles, indicating disrupted cytostome function. Transcriptomic analyses suggested that this deficiency impacted DNA replication and cell cycle regulation. Mass spectrometry-based enzyme assay and lipidomic analysis demonstrated that recombinant PfLPLAT1 exhibited lysophospholipid acyltransferase activity with a preference for unsaturated fatty acids as its acyl donors and lysophosphatidic acids as an acceptor, with its conditional knockout leading to abnormal lipid composition and marked morphological and developmental changes including stage arrest. These findings highlight PfLPLAT1 as a potential target for antimalarial therapy, particularly due to its unique role and divergence from human orthologs.

## Introduction

Malaria is a major mosquito-borne infectious disease that causes a large number of deaths worldwide. During its intraerythrocytic stages, the parasite invades human erythrocytes and encircles itself with a multi-layered membrane structure, which allows communication with the external environment through the uptake of nutrients and the expression of parasitic proteins on the surface of erythrocyte membranes. Both liver-stage and erythrocytic-stage malaria parasites undergo rapid cell proliferation inside the host cells, which requires a large amount of lipid molecules to support the significant increase in the total cell surface area caused by the complex membrane morphology^1^. Therefore, tight regulation of lipid levels and classes are important for the normal proliferation of parasites.

In recent years, lipid metabolism and related areas in malaria parasites have been drawing attention in the study of the roles of lipids in the complex regulation of molecular pathways in parasites and as new drug targets^2^. These studies suggested that parasites can acquire a wide variety of lipids from the erythrocyte cytoplasm and extracellular environment, but they also synthesize them to meet stage-dependent demands for specific lipid classes^3^. Lipid synthesis throughout the life cycle of malaria parasites range from fatty acid synthesis to lipid droplet formation with neutral lipids^2,4^. Although most lipid classes can also be scavenged from the host cytoplasm and extracellular environment, parasites must maintain a stable balance of fatty acids and phospholipid and neutral lipid synthesis to meet the diverse needs of these molecules at different stages and environments^5–8^.

In mammalian cells, phospholipid *de novo* synthesis starts from the incorporation of acyl-CoA into glycerol 3-phophate (G3P) by glycerol 3-phosphate acyltransferases (GPATs) to produce lysophosphatidic acid (LPA). This reaction is followed by additional acyl chain incorporation to produce phosphatidic acid (PA) and subsequent head group modifications to produce diacylglycerol (DAG) and phospholipids (Kennedy cycle)^9^. Acyl chains of these phospholipids are often excised by phospholipases, and the resulting lysophospholipids (LPLs) can be processed back to diacyl phospholipids with different combinations of acyl chains by lysophospholipid acyltransferases (LPLATs) (Lands’ cycle)^10^. To date, 14 mammalian LPLATs have been identified from two independent families, 1-acylglycerol-3-phosphate O-acyltransferase (AGPAT) and membrane-bound O-acyltransferase (MBOAT), based on their primary structures (Table 1). These two families possess AGPAT^11,12^ and MBOAT motifs^13^, respectively, which are essential for LPLAT activity. Each LPLAT has multiple substrate specificities^14^.The study of phospholipid heterogeneity created by a series of acyltransferases and their relative expression levels in tissues has been a rapidly growing area in the understanding of the complex regulation of this cell biology, as well as in the search for unique disease markers.^15,16^

**Table 1.** Currently known human LPLATs.

| Family | Proposed Name | NCBI information |  |  |  |
| --- | --- | --- | --- | --- | --- |
|  |  | Current Official Symbol | Organism | Reference Sequence | Gene ID |
| AGPAT | LPLAT1 | AGPAT1 | human | NM_032741 | 10554 |
|  | LPLAT2 | AGPAT2 | human | NM_006412 | 10555 |
|  | LPLAT3 | AGPAT3 | human | NM_001037553 | 56894 |
|  | LPLAT4 | AGPAT4 | human | NM_020133 | 56895 |
|  | LPLAT5 | AGPAT5 | human | NM_018361 | 55326 |
|  | LPLAT6 | LCLAT1 | human | NM_182551 | 253558 |
|  | LPLAT7 | LPGAT1 | human | NM_014873 | 9926 |
|  | LPLAT8 | LPCAT1 | human | NM_024830 | 79888 |
|  | LPLAT9 | LPCAT2 | human | NM_017839 | 54947 |
|  | LPLAT10 | LPCAT4 | human | NM_153613 | 254531 |
| MBOAT | LPLAT11 | MBOAT7 | human | NM_024298 | 79143 |
|  | LPLAT12 | LPCAT3 | human | NM_005768 | 10162 |
|  | LPLAT13 | MBOAT2 | human | NM_138799 | 129642 |
|  | LPLAT14 | MBOAT1 | human | NM_001080480 | 154141 |

LCLAT: Lysocardiolipin Acyltransferase

In contrast to the mammalian studies, data regarding the equivalent enzymes and the uniqueness of phospholipid synthesis in *Plasmodium* spp. remain limited. Several genes have been suggested as potential acyltransferases by the PlasmoDB database (https://plasmodb.org/plasmo/app/), and only putative GPAT^17–19^ and diacylglycerol acyltransferase (DGAT)^20,21–22^ have been examined for their activity and localization. The ability of GPAT and DGAT to incorporate acyl chains into their substrates, G3P and DAG, respectively, was estimated using whole-cell lysates or by heterogenous expression in *Escherichia coli* (*E. coli*) or yeast, but the enzymatic activity of recombinant parasite proteins was not investigated, probably because of the technical complexity of assessing acyltransferase activity in vitro. Moreover, no studies have been conducted to identify and characterize LPLATs in *P. falciparum*, although two estimated LPLATs were researched in the liver stage of *Plasmodium yoelii*^18^. We performed a database search with the mammalian AGPAT motif (Fig. 1B), which contains four core functional domains^11,23,12^, as a model sequence, and identified a gene in *P. falciparum* that potentially has LPLAT activity: no genes were identified by the MBOAT motif.

**Figure 1.**
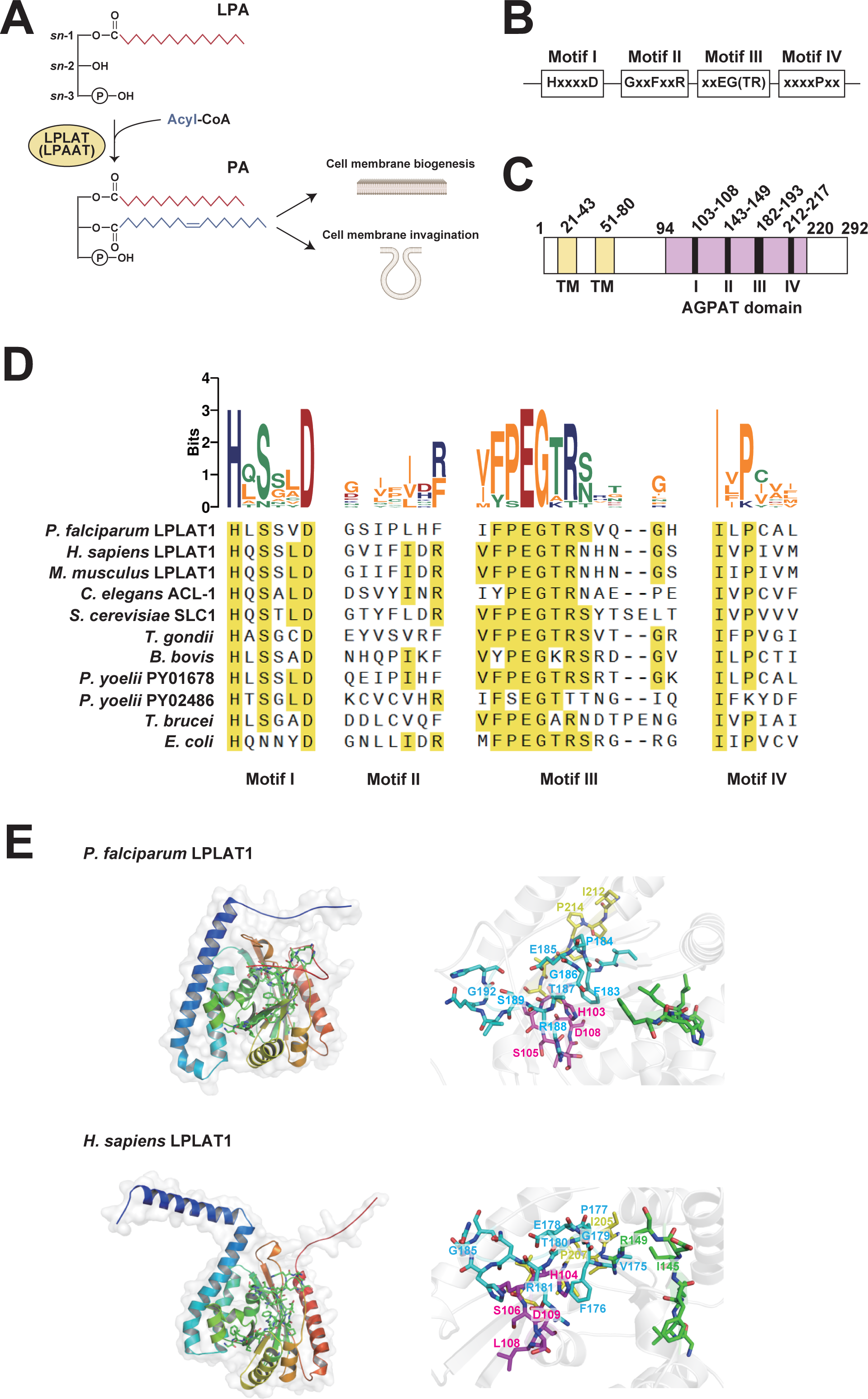
A variety of microorganisms including *P. falciparum* have LPLATs with partially conserved catalytic motifs. (A) Schematic diagram of phosphatidic acid (PA) synthesis from lysophosphatidic acid (LPA). (B) Domain structure of PfLPLAT1. Transmembrane domain (TM) and AGPAT domains (I to IV) were predicted using InterPro. (B) Multiple alignment of catalytic motifs in LPLATs. Amino acid residues with ≥ 50% preservation across described organisms are highlighted in yellow boxes. Acidic, basic, polar, and nonpolar amino acids are shown as red, blue, green, and yellow letters, respectively, in the upper sequence logo. (C) Predicted three-dimensional structures of PfLPLAT1 and HsAGPAT1. Structural data were obtained from the AlphaFold Protein Structure Database. Left panel: overviews. Right panel: magnified views around motifs I–IV. Amino acid residues in motif I-IV are colored in magenta, green, cyan, and yellow. More than 50% preserved residues are labeled.

Here we report the identification and first systematic characterization of a *P. falciparum* LPLAT, which we named PfLPLAT1. Liquid chromatography-tandem mass spectrometry (LC-MS/MS) analysis showed that recombinant PfLPLAT1 incorporates acyl chains into PA (lysophosphatidic acid acyl transferase, or LPAAT activity) and lysophosphatidylcholine (LPC) (lysophosphatidylcholine acyltransferase, or LPCAT activity), with a preference against PA as a substrate (acceptor) and unsaturated fatty acids as acyl chain donors. Conditional knockout of this gene showed a marked growth defect with a failure in stage transition. These phenotypes are also associated with a characteristic cytostome structural abnormality, which is an essential membrane invagination process for the uptake of host erythrocyte cytoplasm to acquire hemoglobin and other nutrients. Our data suggest the importance of PA in cytostome formation and cell cycle progression in malaria parasites.

## Results

### Catalytic motifs of LPLATs are conserved in a wide variety of organisms including *P. falciparum*

In the phospholipid biosynthesis pathway, LPLATs convert LPLs (mono-acyl form) to PLs (di-acyl form) by incorporating an acyl chain into the glycerol back bone (Fig 1A). Mammalian LPLATs are members of either the AGPAT family or MBOAT family, and some mammalian LPLAT isoforms have lysophosphatidylcholine acyltransferase (LPCAT), lysophosphatidylethanolamine acyltransferase (LPEAT), lysophosphatidylinositol aclytransferase (LPIAT), and LPAAT activity in the Lands cycle^14^. Because of the flexible substrate specificity of LPLATs, it is more appropriate to avoid specifying substrate preferences in their names, and simply sort them with numbers^14^.

To identify LPLATs in *P. falciparum*, we first performed a database search using AGPAT motifs^11,12^ of murine LPLAT8 as bait sequences and found an entry designated as a putative *P. falciparum* LPLAT (ID: PF3D7_1444300, designated as 1-acyl-*sn*-glycerol-3-phosphate acyltransferase or LPAAT). We named this protein as *<u>P</u>. falciparum* lyso<u>p</u>hospholipid <u>a</u>cyltransferase <u>1</u> PfLPLAT1, in accordance with previously proposed nomenclature^14^. PfLPLAT1 is predicted to have two transmembrane domains (amino acids 21–43 and 51–80) at the N-terminus and an AGPAT domain (amino acids 94–220) at the C-terminus (Fig 1C). It has been revealed that AGPAT motifs (motifs I–IV) are essential for LPLAT activity in mammalian LPLATs^11,12^. We compared the amino acid sequences with those of putative LPLATs from other organisms (Fig 1D). Multiple alignment showed that motif I (HxxxxD) and motif III (xxEG(TR)) were conserved in *P. falciparum*, while motif II (GxxFxxR) had lower sequence similarity. Motif IV (xxxxPxx) is characterized by a proline residue surrounded by nonpolar amino acids. Although the proline is conserved in PfLPLAT1 motif IV, it is followed by cysteine, which is a polar residue. Considering motif IV is generally less conserved among all AGPATs, it is questionable whether PfLPLAT1 motif IV is involved in the acyltransferase activity. Despite the low conservation of amino acid residues in motif II and IV, prediction of its three-dimensional conformation showed that spatial arrangements of motifs I–IV of PfLPLAT1 were similar to those of *Homo sapiens* LPLAT1 (HsAGPAT1) ^26^ (Fig 1E).

### Recombinant PfLPLAT1 exhibits lysophospholipid acyltransferase activity

Because the AGPAT motifs in PfLPLAT1 are not perfectly conserved, the enzymatic activity of PfLPLAT1 as an acyltransferase must be examined. To confirm whether PfLPLAT1 is an LPLAT, we tested its enzymatic activity by LC-MS/MS analysis, with recombinant PfLPLAT1 synthesized by a wheat germ expression system (Fig 2A). We expressed PfLPLAT1 in proteoliposomes to achieve high purity and to avoid background signals from wheat-germ expression reagents. Both Coomassie Brilliant Blue (CBB) staining and western blotting detected a 35 kDa band for PfLPLAT1 and a faint band at higher molecular weight (approximately 55 kDa), which might be dimerized PfLPLAT1 or complexed with other proteins (Fig. 2B). After purification, PfLPLAT1 was reacted with 6 acyl-CoAs (16:0-, 18:0-, 18:2-, 20:4-, 20:5- and 22:6-CoA) and each deuterium-labeled LPL (d9-LPA(16:0), d31-LPC(16:0), or d9-LPE(16:0)) to produce diacyl phospholipid products PAs, PC, and PEs). Deuterated LPLs enabled us to distinguish newly synthesized lipids from residual lipids of the same type. In a given substrate-rich condition, six deuterated PAs, four PCs, and faint activity for PE were detected (Fig. 2C, D, and E). In all PA products, PA(34:2) and PA(36:5) were the most abundant, showing that PfLPLAT1 utilized linoleic acids (18:2-CoA) and eicosapentaenoic acids (20:5-CoA) as acyl chain donors. These acyl-CoA moieties were also used and produced PC(34:2) and PC(36:5). As for saturated and monounsaturated lipids, PfLPLAT1 used16:0-CoA and 18:1-CoA as donors for both LPA and LPC to a slightly lower extent. Taken together, this enzyme assay revealed that PfLPLAT1 could utilize LPAs and LPCs as its substrates and may prefer to use polyunsaturated fatty acids in the PA production.

**Figure 2.**
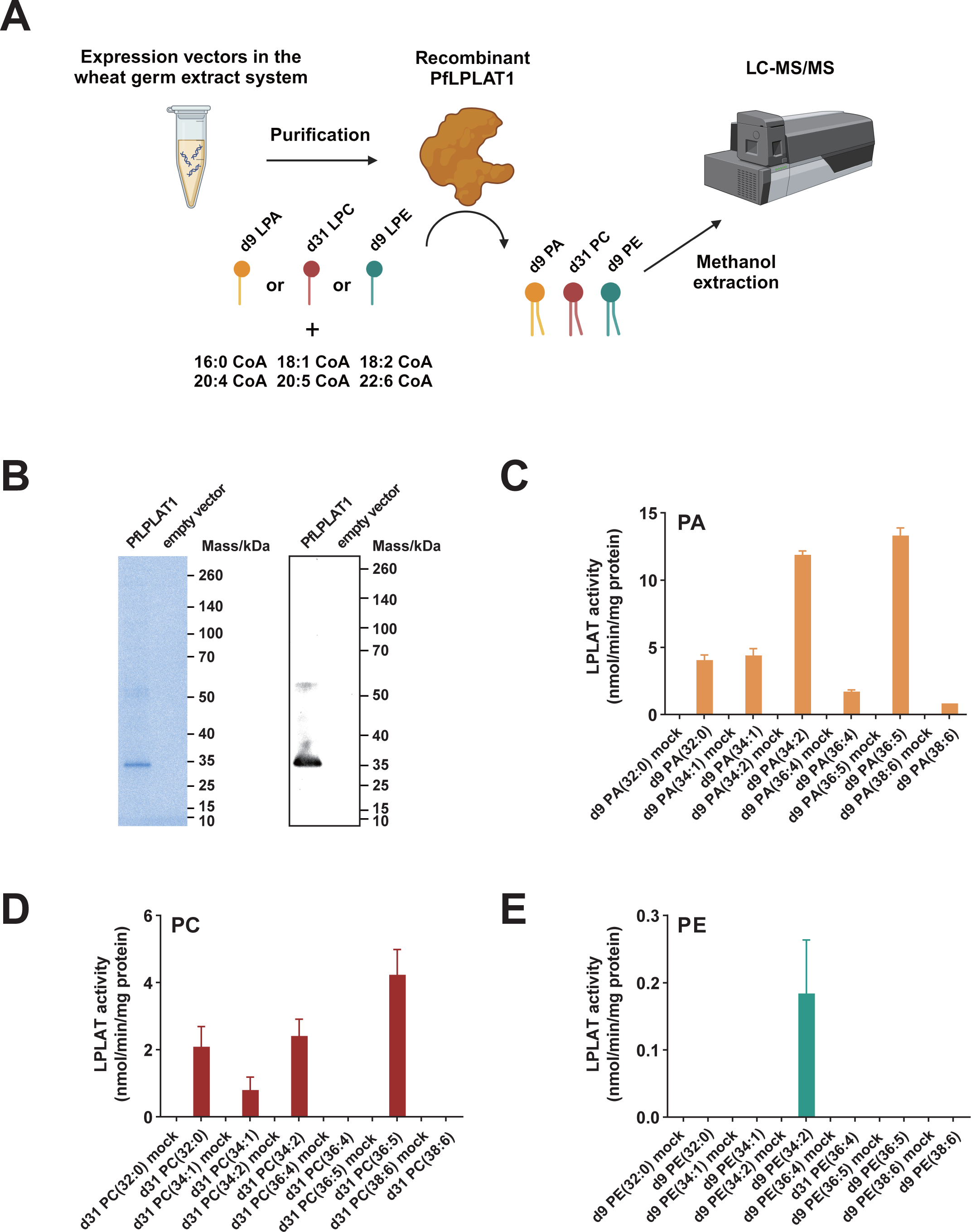
PfLPLAT1 shows LPAAT and LPCAT enzymatic activity. (A) Schematic image of LC-MS/MS based in vitro enzyme assay. Mixture of fatty acids (acyl chain donors) are reacted with a deuterium-labeled LPLs in the presence of PfLPLAT1 and diacyl forms of each lipid type with deuterium labeling were quantified. (B, C) Recombinant PfLPLAT1 in proteo-liposomes generated from the wheat germ expression system was detected by western blotting analysis (B) and Coomassie Brilliant Blue (CBB) staining at the theoretical size (35 kDa). (D, E, F) PA (D), PC (E), and PE (F) were estimated by LC-MS/MS-based analysis based on signal intensities from standards (PA(34;1) for PA, PC(34:1) for PC, and PE(34:1) for PE estimation). The x-axis shows product diacyl phospholipids with the sum of the carbons and double bonds in two acyl chains (i.e., PA(34:1): total number of carbons in acyl chains = 34 and the number of double bonds = 1.

### PfLPLAT1 is essential in asexual replication and stage transition of *P. falciparum*

To study the role of *Pflplat1* in the asexual life cycle of parasites, we generated conditional knockout parasite lines, which were referred to as *Pflplat1*:LoxPint:HA, using the dimerizable Cre recombinase (DiCre) system^27,28^. We incorporated the plasmid DNA targeting native *Pflplat1* locus with the selection-linked integration (SLI) method^29^ (Fig 3A) and confirmed the integration by a diagnostic PCR (Fig 3B and Fig S1A). To check whether the conditional knockout system works in established parasite lines, PCR, western blotting, and indirect immunofluorescence assay (IFA) were performed (Fig 3C–E and Fig S1B–D). Shifts in DNA bands were observed in rapamycin-treated *Pflplat1*:LoxPint:HA C9 and G6 clonal lines, which indicated that *Pflplat1* locus flanked by LoxP sites was excised and shortened (Fig 3C and Fig S1B). Western blotting analysis revealed decreased protein expression of PfLPLAT1 after 24 hours (fractional decrease in band density: 72.2% ± 11.2 (C9), 59.8% ± 25.1 (G6)) or 48 hours (95.9 ± 3.12 (C9), 91.3 ± 4.89 (G6)) of rapamycin treatment (Fig 3D and Fig S1C). Moreover, IFA showed a significant decrease in the immunofluorescence intensity of PfLPLAT1 after 24 hours of rapamycin treatment (Fig 3E and Fig S1D). Taken together, *Pflplat1* was efficiently knocked out in transgenic parasites in the presence of rapamycin.

**Figure 3.**
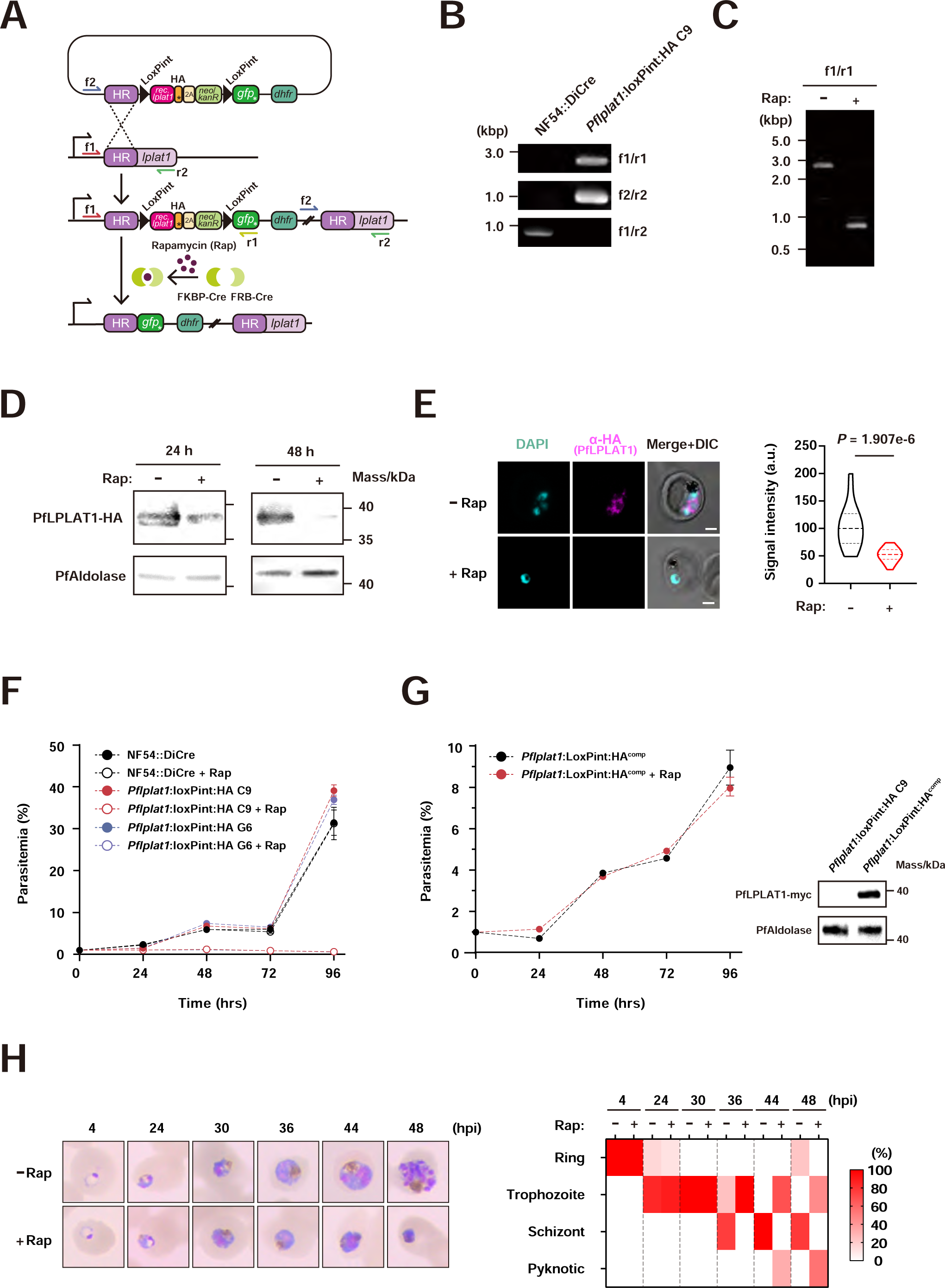
PfLPLAT1 is essential in asexual replication and stage transition of *P. falciparum*. (A) Conditional gene knockout design for *Pflplat1* with the DiCre system. Arrows indicate a transcription initiation site and the primer binding sites for PCR-based tests. (B) Diagnostic PCR showed that the target gene was successfully replaced with the targeting constructs: DNA segments were correctly amplified with f1/r1 (NF54::DiCre, not amplified; *Pflplat1*:LoxPint:HA C9, 2676 bp), f2/r2 (NF54::DiCre, not amplified; *Pflplat1*:LoxPint:HA C9, 979 bp), and f1/r2 (NF54::DiCre, 1027 bp; *Pflplat1*:LoxPint:HA C9, not amplified) primer sets. (C–E) Conditional gene disruption of *Pflplat1* was confirmed by PCR, western blotting, and immunofluorescence assay (IFA). (C) The size of DNA amplicons for the 5’ segment was shifted from 2676 to 784 bp at 24 hours of rapamycin treatment. (D) Reductions in PfLPLAT1 were detected in *Pflplat1*:LoxPint:HA clone C9 with anti-HA antibody at 24 and 48 hours of rapamycin treatment. PfAldolase was used as an internal control for the total amount of proteins. Theoretical molecular mass/kDa of HA-tagged PfLPLAT1 and PfAldolase is 37.4 and 40.0, respectively. Experiments were repeated three times. (E) Left panel: representative immunofluorescence images of *Pflplat1*:LoxPint:HA C9 at 24 hours of rapamycin treatment. Samples were stained for nuclei with DAPI (cyan) and PfLPLAT1 with anti-HA antibody (magenta) and analyzed by confocal microscopy. Scale bars: 2 μm. Right panel: violin plots showing the fluorescence intensity of HA-tagged PfLPLAT1 in rapamycin-treated (+Rap) or untreated (−Rap) *Pflplat1*:LoxPint:HA C9 (n = 20 per group). Violin plots range from the minimum to the maximum values. Dotted bottom, middle, and top horizontal lines show 25th percentile, median, and 75th percentile, respectively. Experiments were repeated two times. Statistical analysis used F-test for equality of variance followed by two-tailed t-test. (F) Growth curve of +Rap or −Rap *Pflplat1*:LoxPint:HA C9 and G6 clones. Data are shown as mean ± SD from n = 3 independent biological replicates. (G) Overexpressed PfLPLAT1 rescued parasite growth under the rapamycin treatment. Left panel: growth curve of +Rap or −Rap *Pflplat1*:LoxPint:HA^comp^. Data is shown as mean ± SD from n = 3 independent biological replicates. Right panel: myc-tagged PfLPLAT1 expression in *Pflplat1*:LoxPint:HA^comp^. PfAldolase was used as an internal control for total amount of protein. The theoretical molecular mass/kDa of myc-tagged PfLPLAT1 and PfAldolase is 40.0. (H) Stage distributions of +Rap or −Rap *Pflplat1*:LoxPint:HA C9. Left panel: representative images of Giemsa-stained blood films at described time points (4, 24, 30, 36, 44, and 48 hours post-infection [hpi]). Right: heatmap showing the parasite stage distributions at the described time points. Data is shown as mean from n = 3 independent biological replicates.

To study the effect of *Pflplat1*-defect on the asexual replication of parasites, we performed growth assays using synchronized *Pflplat1*:LoxPint:HA C9 and G6 clonal lines. The parasitemia of rapamycin-treated (+Rap) parasites did not increase and recover until the end of assay, while that of rapamycin-untreated (−Rap) parasites showed normal growth (Fig 3F). This clearly indicated that *Pflplat1* is essential for parasite growth in the erythrocytic stages, which is concordant with the low mutagenesis index score of *Pflplat1* shown in previous genome wide screening^30^. *Pflplat1*:LoxPint:HA C9 clonal line was used in further analyses because no obvious differences in gene knockout efficiency and growth kinetics between two parasite clones were observed in the presence of rapamycin. To test if the growth defect of rapamycin-treated *Pflplat1*:LoxPint:HA was due to the ablation of *Pflplat1,* we established a genetically complementary line, *Pflplat1*:LoxPint:HA^comp^, expressing episomal PfLPLAT1 (Fig 3G). Growth assays of this parasite line showed that +Rap parasites were able to grow to the same extent as rapamycin-nontreated parasites, which indicated that aberrant growth of +Rap *Pflplat1*:LoxPint:HA was due to the lack of PfLPLAT1 (Fig 3G). Analyses of stage distribution of *Pflplat1*:LoxPint:HA with or without rapamycin treatment revealed clear differences in their stage at 36 hours post-infection (hpi) (Fig 3H). −Rap parasites entered into the schizont stage at 36 hpi, while +Rap parasites did not exhibit cell division. The growth of +Rap parasites was stalled in the trophozoite stage when observed at 44 and 48 hpi. Moreover, shrunk and pyknotic parasites were observed in the +Rap group at 44 hpi and approximately 50% of parasites were found in the aberrant form at 48 hpi. This suggests that *Pflplat1* is indispensable for stage transition from trophozoites to schizonts.

Previous transcriptional analyses showed that *Pflplat1* is highly expressed in the gametocyte stage^31^, which encouraged us to investigate the essentiality of *Pflplat1* at this stage. Parasites were treated with rapamycin at day 10 after gametocytogenesis, and gametocytemia was measured on days 12–15 post-gametocytogenesis. Although *Pflplat1* was excised in the presence of rapamycin (Fig S2A), +Rap and −Rap parasites showed no significant difference in gametocytemia and gametocyte stage distribution (Fig S2B, S2C). This suggests that *Pflplat1* is not essential in the maturation process of gametocytes.

### PfLPLAT1 colocalizes with endoplasmic reticulum throughout the asexual stage

It was shown that most enzymes functioning in phospholipid synthesis were integrated into the endoplasmic reticulum (ER) membrane in mammalian cells^32^. To investigate whether PfLPLAT1 localizes on the ER, we performed an indirect IFA co-stained with an ER marker protein BiP (binding immunoglobulin protein) and confocal microscopy. Signal profiles of PfLPLAT1 and BiP showed similar patterns throughout all asexual stages, which suggests that most PfLPLAT1 localizes on the ER (Fig 4, right panel). In early and middle schizont stages, PfLPLAT1 and BiP were highly co-localized based on the Pearson’s r values (Fig 4, right panel). These two stages also showed expression peaks of PfLPLAT1 and BiP^33^. Our results indicated that the ER is a fundamental platform for phospholipid synthesis in *P. falciparum*, and demand for phospholipids by parasites possibly reaches the highest level in the early and middle schizont stages, in which parasites undergo multiple nuclear divisions.

**Figure 4.**
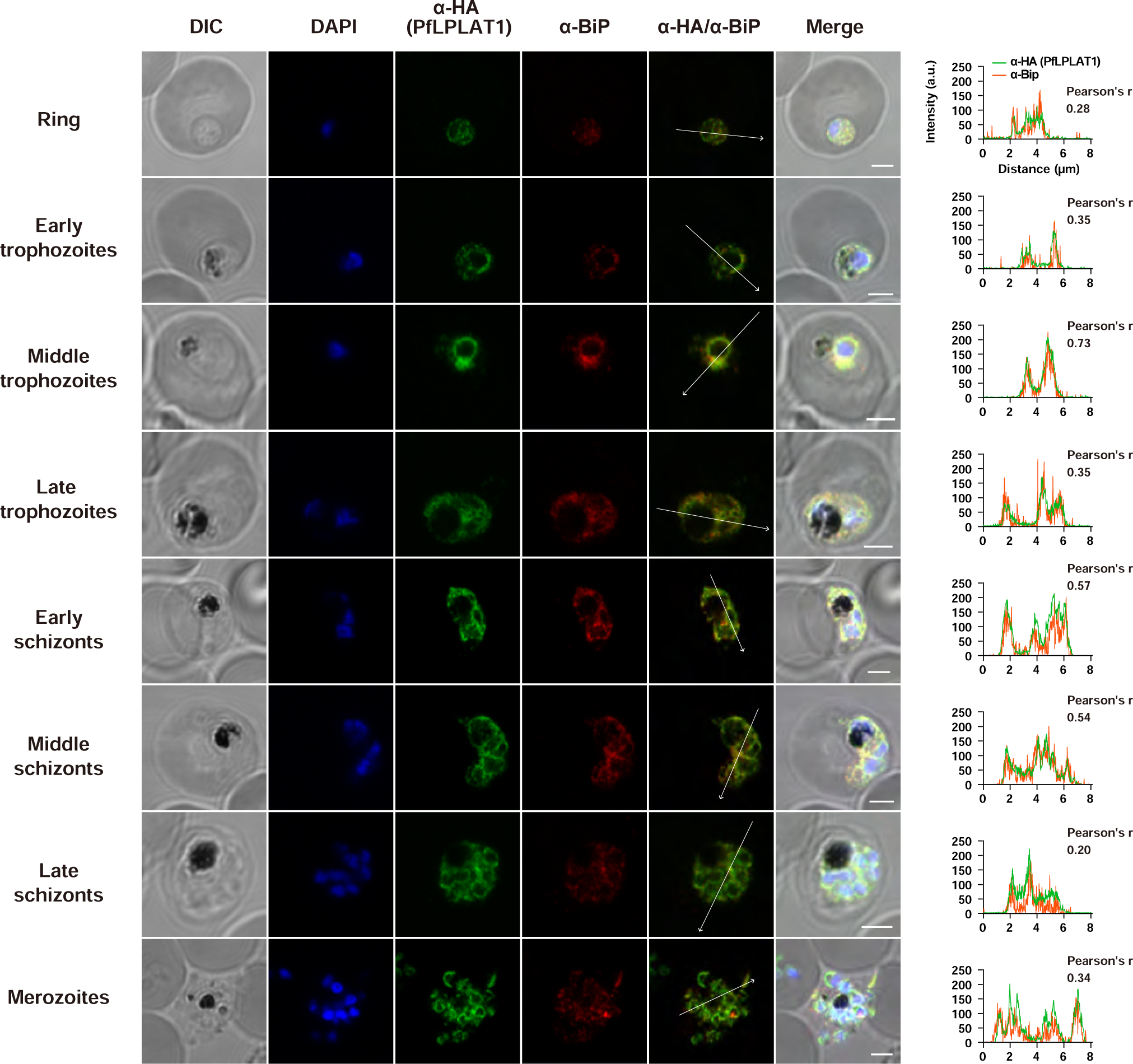
PfLPLAT1 colocalizes with endoplasmic reticulum throughout the all-asexual stage. Left panel: representative confocal microscope images of parasites co-stained with PfLPLAT1 and ER-resident protein BiP. Samples were stained for nuclei with DAPI (blue), PfLPLAT1 with anti-HA antibody (green), and ER with anti-BiP antibody (red). Scale bars = 2 μm. Right panel: signal profiles of anti-HA (green) and anti-BiP (red) on white arrows in the overlays. Pearson’s r values were calculated using overlays of anti-HA and anti-BiP signal intensities to show the overlapping degree of both signals.

### *Pflplat1* deletion caused abnormal cytostome formation and their internalization

Our previous studies showed that mammalian LPLATs maintain the integrity of cellular membranes in various tissues and their functions^15,16^. To study the effect of PfLPLAT1 deletion on the membrane morphology of parasites, we observed parasitized erythrocyte structure at 36 hpi using a high-voltage scanning electron microscope (HV-STEM), which enables sample sections of 250–1,000 nm thickness with 1000 keV accelerating voltage to be obtained^34,35^. Due to the deeper penetration of electrons into the target specimen, HV-STEM provides more representative images of subcellular structures within a space. Sections (250 nm thickness) of rapamycin-treated parasites showed gigantic omega-shaped and open-mouthed vacuoles (Fig 5C-K); such a structure was not observed in −Rap parasites (Fig 5A and B). These vacuoles showed the typical characteristics of cytostomes, which comprise double membranes (Fig 5H) derived from the parasite plasma membranes (PPMs) and parasitophorous vacuole membranes (PVMs) and collars around the opening^36–38^. Although cytostomes with a normal appearance (diameter of approximately 190 nm) were also observed in +Rap samples (Fig 5J)^39^, the outer diameter of abnormal cytostomes reached approximately 500 nm (Fig 5F and K). Furthermore, several vacuoles were observed between PPM and PVM in *Pflplat1*-deficient parasites (Fig 5G), which may be transported from either side of the PPMs and PVMs.

**Figure 5.**
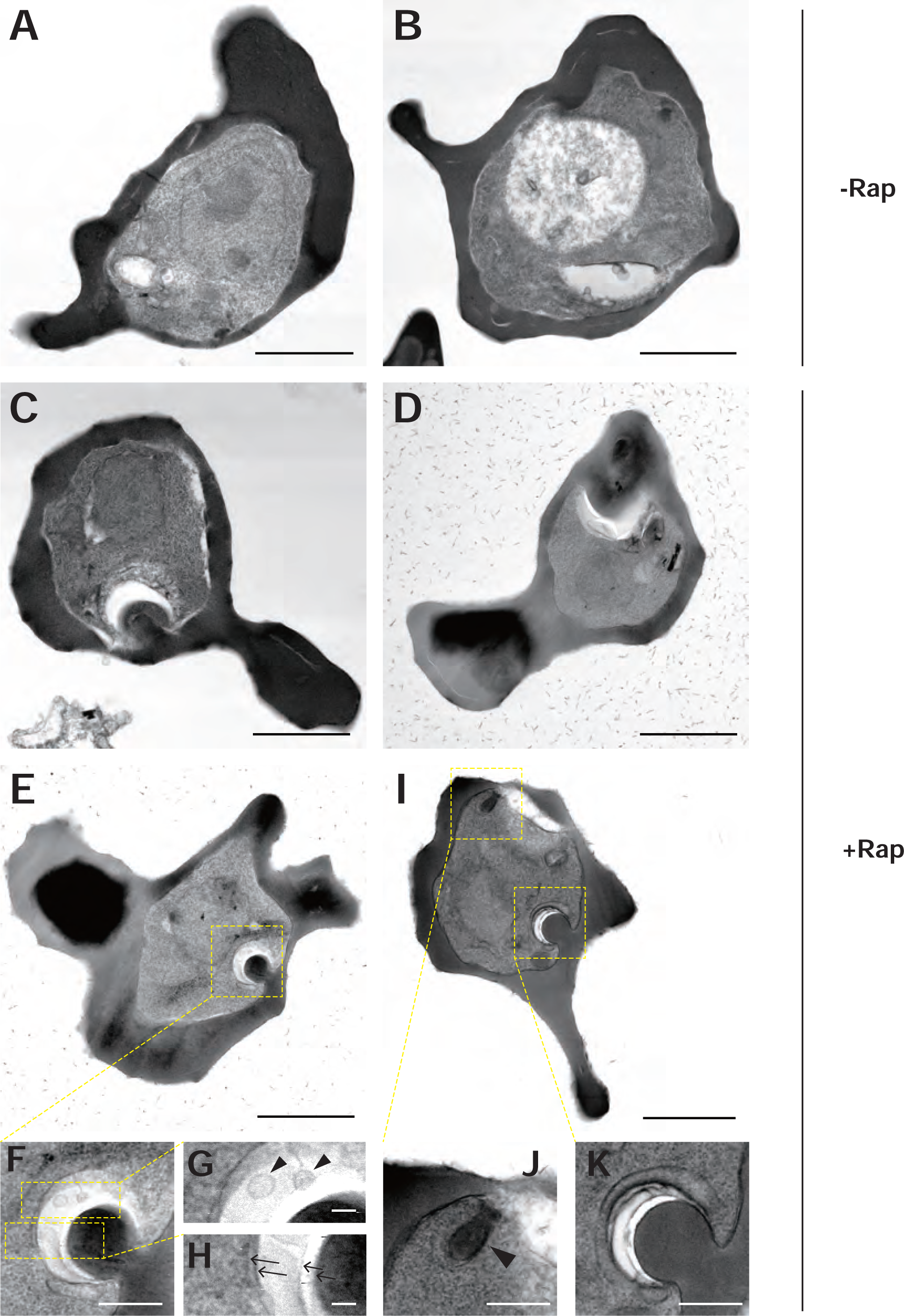
*Pflplat1* deficiency led to the formation of abnormal cytostomes in parasites. STEM images (bright-field) of −Rap (A, B) and +Rap (C-K) *Pflplat1*:LoxPint:HA at 36 hpi with 250 nm sections were obtained by a high-voltage transmission electron microscope. (F) Magnified image of Figure 5E in a yellow dotted rectangle. (G, H) Magnified images of Figure 5F in top (G) and bottom (H) yellow dotted rectangles. Black arrowheads in (G) show vesicle-like structures and black arrows in (H) show the parasite PPM and PVM. (J, K) Magnified images of Figure 5I in upper-left (J) and bottom-right (H) yellow dotted rectangles. Black arrowhead in (J) shows a cytostome with normal appearance. Scale bars = 2 μm (A–E), 500 nm (F, J, K), and 100 nm (G, H). Small needle-like structures around the erythrocytes in Fig 5D and E are agar particles, and they did not influence the internal structures of parasitized erythrocytes.

*P. falciparum* acquires hemoglobin and other nutrients through the cytostome and digests hemoglobin into amino acids in digestive vacuoles. Therefore, we hypothesized that failures of fissions of cytostomes could lead to decreased hemoglobin uptake and reduced size of hemozoins, which are crystalline products of degraded hemoglobin. We quantified the size of hemozoins using polarized light microscopy^40^, which identify hemozoins with a high sensitivity due to the birefringence caused by crystalline arrangements of β-hematin. In rapamycin-treated parasites at 30 hpi, the size of hemozoins was significantly decreased compared with that of rapamycin-nontreated parasites (Fig S3). These findings support our hypothesis and indicate that *Pflplat1*-deficiency leads to an insufficient supply of nutrients and amino acids in parasites through the formation of abnormal cytostomes.

### Transcriptomic profiles reveal decreased gene expression related to DNA replication and cell cycle regulation in *Pflplat1*-deficient parasites

To explore biological mechanisms that cause the death of *Pflplat1*-deficient parasites at the late-trophozoite stage, we studied the transcriptomes of +Rap and −Rap *Pflplat1*:LoxPint:HA lines with samples collected at 24 and 30 hpi: timepoints before the appearance of dying parasites. Although few differentially expressed genes (DEGs) were detected at 24 hpi (Fig S4, Table S2) and the gene expression patterns were not well clustered together in +Rap or −Rap samples (Fig S4), 267 upregulated genes (log_2_FC > 0.5, adjusted p-value < 0.5) and 331 downregulated genes (log_2_FC < 0.5, adjusted p-value < 0.5) were observed at 30 hpi (Fig 6A, Table S2). The gene expression profiles of +Rap or −Rap parasites at 30 hpi were well clustered together, indicating sufficient reproducibility of gene expression analysis at this time point (Fig S4). It was shown that 107 of 267 upregulated genes were classified as *Plasmodium* interspersed repeat genes, almost all of which are highly expressed at the early trophozoite stage^31,33^. This result indicates the possibility that *Pflplat1*-deficiency leads to delayed gene expression in +Rap parasites. Three hundred thirty-one downregulated genes were shown to function in various biological process such as the lipid metabolic pathway, DNA replication, GPI (glycosylphosphatidylinositol) anchor biosynthetic process, metal ion transport, and the ERAD (ER-associated degradation) pathway (Fig 6B and 6C), suggesting that *Pflplat1* deficiency affects a wide variety of metabolic pathways in parasites. Downregulated genes are categorized into DNA replication, cell cycle, DNA metabolic process, cell cycle regulation, and nuclear division, which might explain the growth arrest of *Pflplat1*-deficient parasites at the late trophozoite stage (Fig 3H).

**Figure 6.**
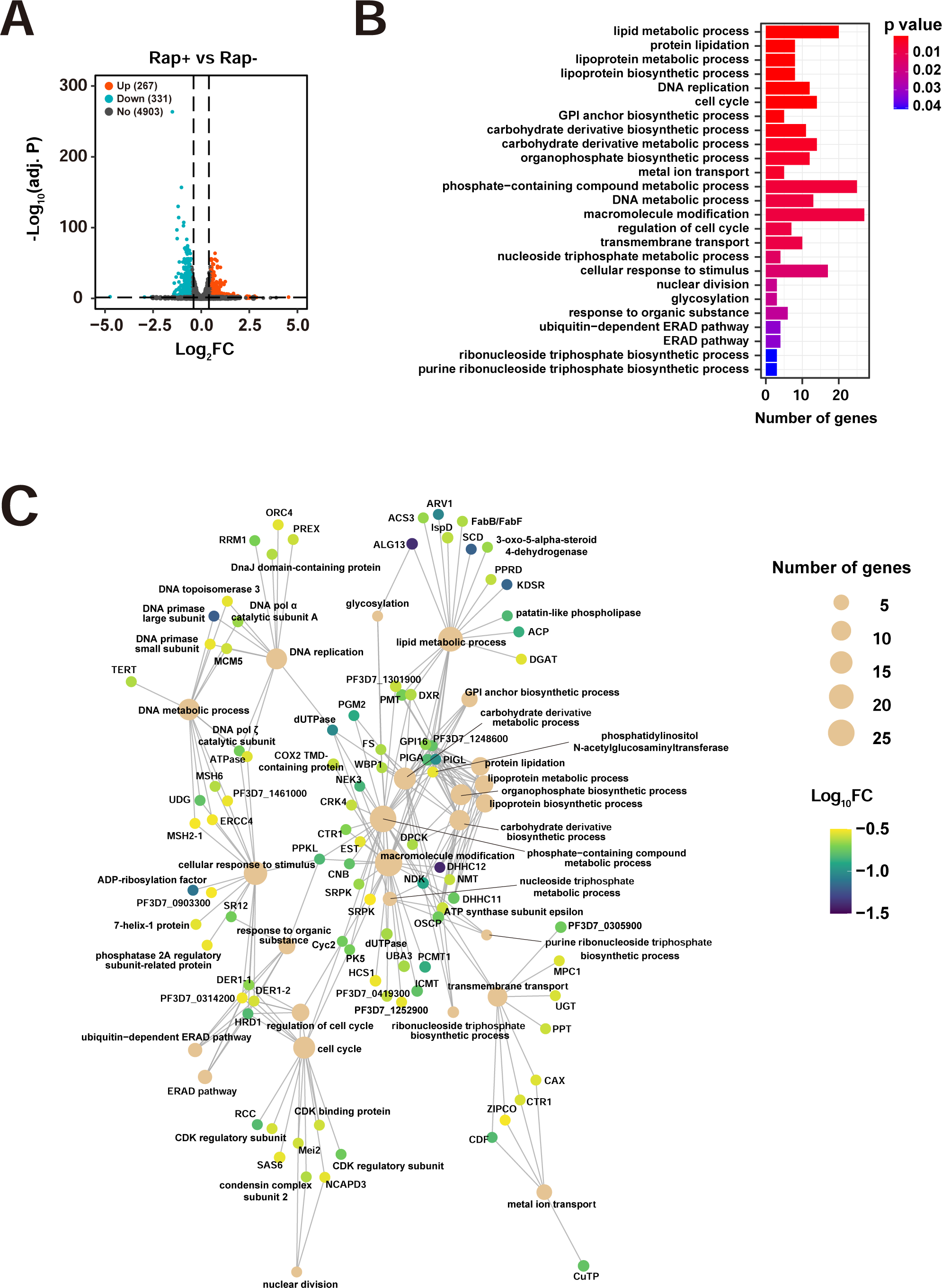
*Pflplat1* deficiency influences the parasite transcriptomic profiles related to cell cycle progression. (A) Volcano plot showing differentially expressed genes in +Rap versus −Rap *Pflplat1*:LoxPint:HA at 30 hpi (n = 3 independent biological replicates per group). Red dots are upregulated genes (log_2_FC > 0.5, adjusted p-value < 0.05) and blue dots are downregulated genes (log_2_FC < 0.5, adjusted p-value < 0.05). Dotted vertical lines show 0.5 and −0.5 of log_2_FC and dotted horizontal line shows 0.05 of adjusted p-value. Adjusted p-values were calculated using the Benjamini–Hochberg method. (B) Bar chart showing the number of downregulated genes classified into each biological process. P-values were calculated using the hypergeometric test. (C) Network diagram showing linkage of downregulated genes that are categorized in described biological processes. ACP, acyl carrier protein; ACS3, acyl-CoA synthetase 3; CAX, cation/H+ antiporter; CDF, cation diffusion facilitator; CNB, calcineurin subunit B; CRK4, cdc2-related protein kinase 4; CTR1, copper transporter 1; CuTP, copper-transporting ATPase; DGAT, diacylglycerol O-acyltransferase; DPCK, dephospho-CoA kinase; dUTPase, dUTP pyrophosphatase; DXR, 1-deoxy-D-xylulose 5-phosphate reductoisomerase; EST, exported serine/threonine protein kinase; FabB/FabF, 3-oxoacyl-acyl-carrier protein synthase I/II; FS, GDP-L-fucose synthase; GPI16, GPI transamidase component GPI16, putative; HCS1, biotin-protein ligase 1; ICMT, protein-S-isoprenylcysteine O-methyltransferase; IspD, 2-C-methyl-D-erythritol 4-phosphate cytidylyltransferase; KDSR, 3-ketodihydrosphingosine reductase; MCM5, minichromosome maintenance complex component 5; Mei2, MEI2-like RNA-binding protein; MPC1, mitochondrial pyruvate carrier protein 1; MSH2-1, MutS protein homolog 2; MSH6, mutS homolog 6; NCAPD3, condensin-2 complex subunit D3; NDK, nucleoside diphosphate kinase; NEK3, NIMA-related kinase 3; NMT, peptide N-myristoyltransferase; ORC4, origin recognition complex subunit 4; OSCP, ATP synthase subunit O; PCMT1, protein-L-isoaspartate(D-aspartate) O-methyltransferase 1; PGM2, phosphoglucomutase-2; PIGA, phosphatidylinositol N-acetylglucosaminyltransferase subunit A; PIGL, N-acetylglucosaminyl-phosphatidylinositol de-N-acetylase; PK5, protein kinase 5; PMT, phosphoethanolamine N-methyltransferase; PPKL, protein phosphatase-containing kelch-like domains; PPRD, polyprenol reductase; PREX, plastid replication-repair enzyme; PPT, phosphoenolpyruvate/phosphate translocator; RCC, regulator of chromosome condensation; RRM1, ribonucleotide reductase catalytic subunit M1; SAS6, spindle assembly abnormal protein 6; SCD, stearoyl-CoA desaturase; SR12, serpentine receptor 12; SRD5A, 3-oxo-5-alpha-steroid 4-dehydrogenase; SRPK, serine/threonine protein kinase; TERT, telomerase reverse transcriptase; TOP3, DNA topoisomerase 3; UBA3, ubiquitin like modifier activating enzyme 3; UDG, uracil-DNA glycosylase; UGT, UDP-galactose transporter; ZIPCO, ZIP domain-containing protein.

### Lipidomic analysis reveals the effect of PfLPLAT1 on lipid profiles of malaria parasites

To investigate the effect of PfLPLAT1 deletion on the profile of phospholipid species in parasites, we performed a LC-MS/MS-based lipidomic analysis using +Rap and −Rap *Pflplat1*:LoxPint:HA. Phospholipids were extracted from parasites at 30 hpi and subjected to LC-MS/MS. In +Rap parasites, 9 of 10 PAs, 4 of 14 Pes, and 16 of 24 PCs but only 2 of 12 sphingomyelies (SMs) species were decreased compared with those in −Rap parasites (Fig 7A–D). The sum of PA, PE, and PC decreased in +Rap parasites, while that of SM did not change (Fig 7E). This indicates that PfLPLAT1 may also have activity for producing diacyl forms of PA, PE, and PC in parasites, which is partly consistent with the *in vitro* enzymatic activity of PfLPLAT1 (Fig 2C–E). Furthermore, the relative amount of PA(32:0), PA(34:0), PA(34:1), PC(36:5), and SM(42:3) was decreased, whereas that of PA(34:2) and PC(31:0) was increased in +Rap parasites (Fig S5). Collectively, *Pflplat1* deficiency led to decreased amounts of phospholipids and affected these phospholipid compositions in parasites.

**Figure 7.**
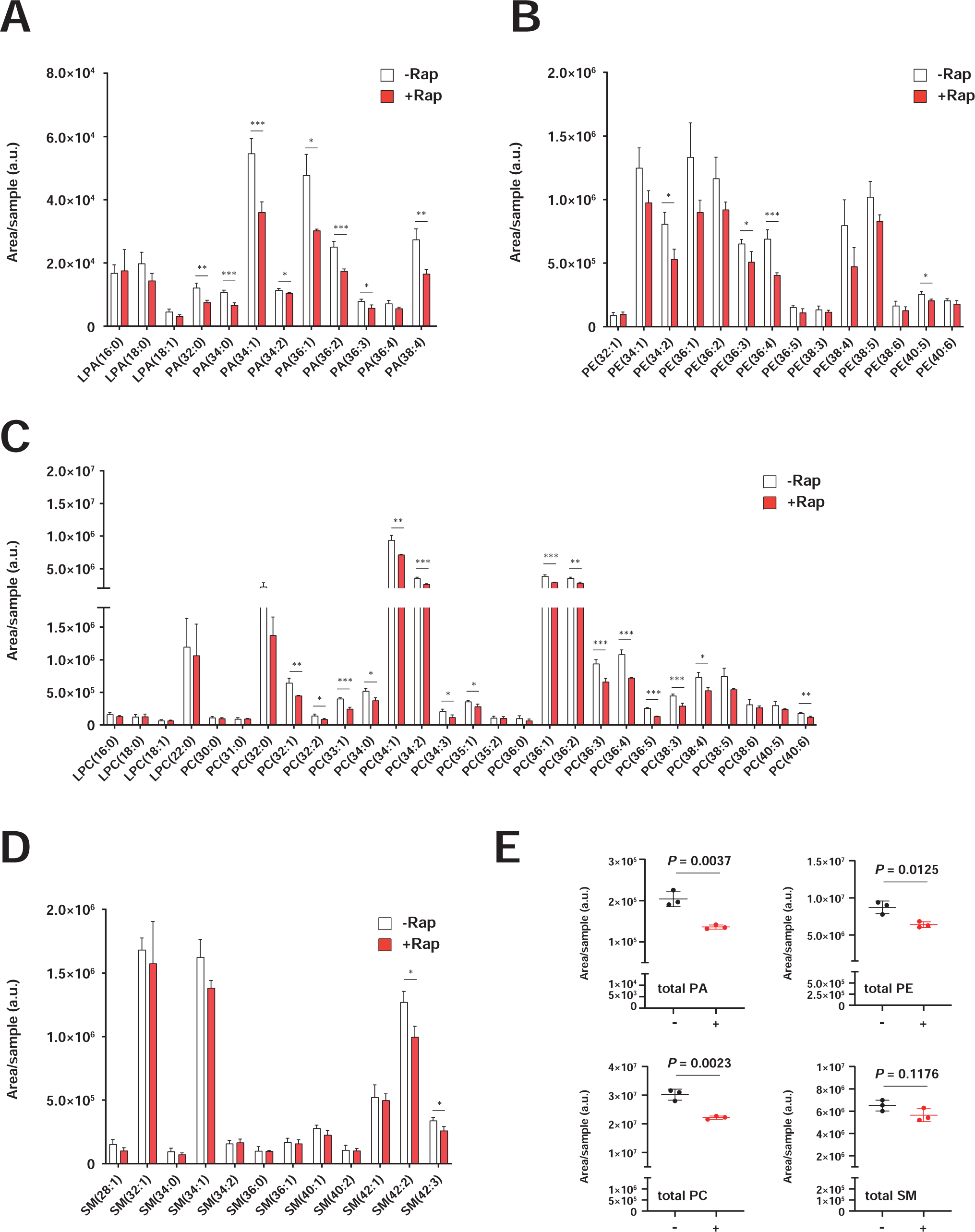
*Pflplat1* deficiency induced modified phospholipid profiles in parasites. (A–D) Bar graphs showing area values of PA (A), PE (B), PC (C), and SM species (D) in +Rap and −Rap *Pflplat1*:LoxPint:HA at 30 hpi. Extracted lipids from an equal number of parasites were analyzed by LC-MS/MS. (E) Dot plots showing the total area values of PA (excluding LPA signals), PE, PC (excluding LPC signals), and SM in −Rap and +Rap *Pflplat1*:LoxPint:HA at 30 hpi. Data are shown as mean ± SD from n = 3 independent biological replicates. P-values were calculated using unpaired two-tailed t-test (*, *P* < 0.05; **, *P* < 0.01, ***, *P* < 0.005).

## Discussion

Phospholipids are *de novo* synthesized in the Kennedy cycle,^9^ and acyl chains of these phospholipids can be exchanged by phospholipase A2 and LPLATs in the Lands cycle^10^. This process maintains phospholipid diversity in tissues for their function. More than 14 LPLATs are known to function in these cycles, and several LPLs are frequently used as substrates by a single LPLAT^14^. In our enzyme assay, recombinant PfLPLAT1 expressed by a wheat germ *in vitro* expression system showed high specificity against LPAs and a mild preference for LPCs as acyl chain acceptors. This finding highlighted that PfLPLAT1 functions not only as an LPAAT but also as an LPCAT, showing that *P. falciparum* may also undergo the Lands cycle pathway. A previous bioinformatic analysis suggested that there are 26 phospholipases encoded by the parasite’s genome^41,42^, while PfLPLAT1 described in this report is one of a few entries we identified from our database search. In contrast to the multiple LPLATs and phospholipases involved in the Lands cycle, *P. falciparum* may have a few LPLATs, which in turn suggests that PfLPLATs may be critical for the regulation of phospholipid biosynthesis. We are still characterizing other putative PfLPLATs regarding their enzymatic activities, but, considering the limited possibility of PfLPLATs and that PfLPLAT1 is an essential enzyme, parasite LPLATs might have activity against multiple LPL species in addition to LPA and LPC to control the optimal profile of lipid species for parasite cell homeostasis.

PfLPLAT1 preferentially uses unsaturated fatty acids such as linoleic acids and eicosapentaenoic acids as substrates (Fig 2C and D). Lipids with unsaturated fatty acids have larger molecular cross-sectional areas that may ease molecular packing and increase membrane flexibility. This indicates that PfLPLAT1 helps the parasite plasma membrane to maintain a high flexibility and/or low viscosity to deal with acute deformations of the parasite membrane during its rapid replication.

While fatty acid synthesis, neutral lipid storage, cholesterol transport, and phospholipase studies in parasites have progressed, development of a comprehensive understanding of the phospholipid synthesis pathway remains insufficient. Several transgenic parasites to conditionally target essential genes related to lipid metabolism have been established, and their growth kinetics were studied upon induction of gene deletion^41,43–46^. However, the growth defects of gene-deficient parasites were observed only after several cycles had passed, in contrast to the obvious growth abnormality detected in *Pflplat1*-deficient parasites at the first cycle. This result indicates that the PfLPLAT itself and/or downstream of lipid maintenance in the parasite is highly important for their normal growth.

Abnormal cytostomes in *Pflplat1*-deficient parasites occurred during the course of parasite death, which has not been reported in previous studies. Cytostomes occur in a clathrin-independent manner in *P. falciparum*,^47^ and dynamin-like protein 1 (PfDYN1) does not localize on the collars of cytostomes^48^. Moreover, PfDYN1 and PfDYN2 were not identified as downregulated genes in our transcriptomic analysis (Fig. 6), suggesting that PfDYNs are likely independent of the formation of abnormal cytostomes, regardless of their ability to pinch off the membrane. Our EM data showed that abnormal cytostomes had large openings that were at least three times the diameter of normal cytostomes. This indicates that the physical states of the openings with negative membrane curvature are different from those of normal cytostomes. PA and PE have molecular shapes of inverted cones and generate negative curvature, which leads to efficient vesicle fusions, fissions^49,50^, and cytokinesis in mammalian cells^51^. Considering these physical properties of PA, as well as the reduced levels of PA, PE, and PC, it is reasonable to speculate that lipid profile changes in *Pflplat1* deficiency could interrupt cytostome fission and lead to the formation of gigantic vesicles due to PA shortage in the cell membrane (Fig. 8). Downregulation of *P. falciparum* homolog of vesicle-associated membrane protein 8 (VAMP8), a SNARE protein on lysosomes, in *Pflplat1*-deficient parasites also may drive further abnormalities. VAMP8 regulates lysosome–autophagosome associations^52,53^ and SNARE proteins are proposed to function in cytostome-digestive vacuole (DV) fusion^48^. Thus, dysregulation of cell membrane curvature and fusion of cytostomes with DVs by downregulated VAMP8 possibly leads to abnormal hemoglobin digestion in a coordinated manner (Fig. 8).

**Figure 8.**
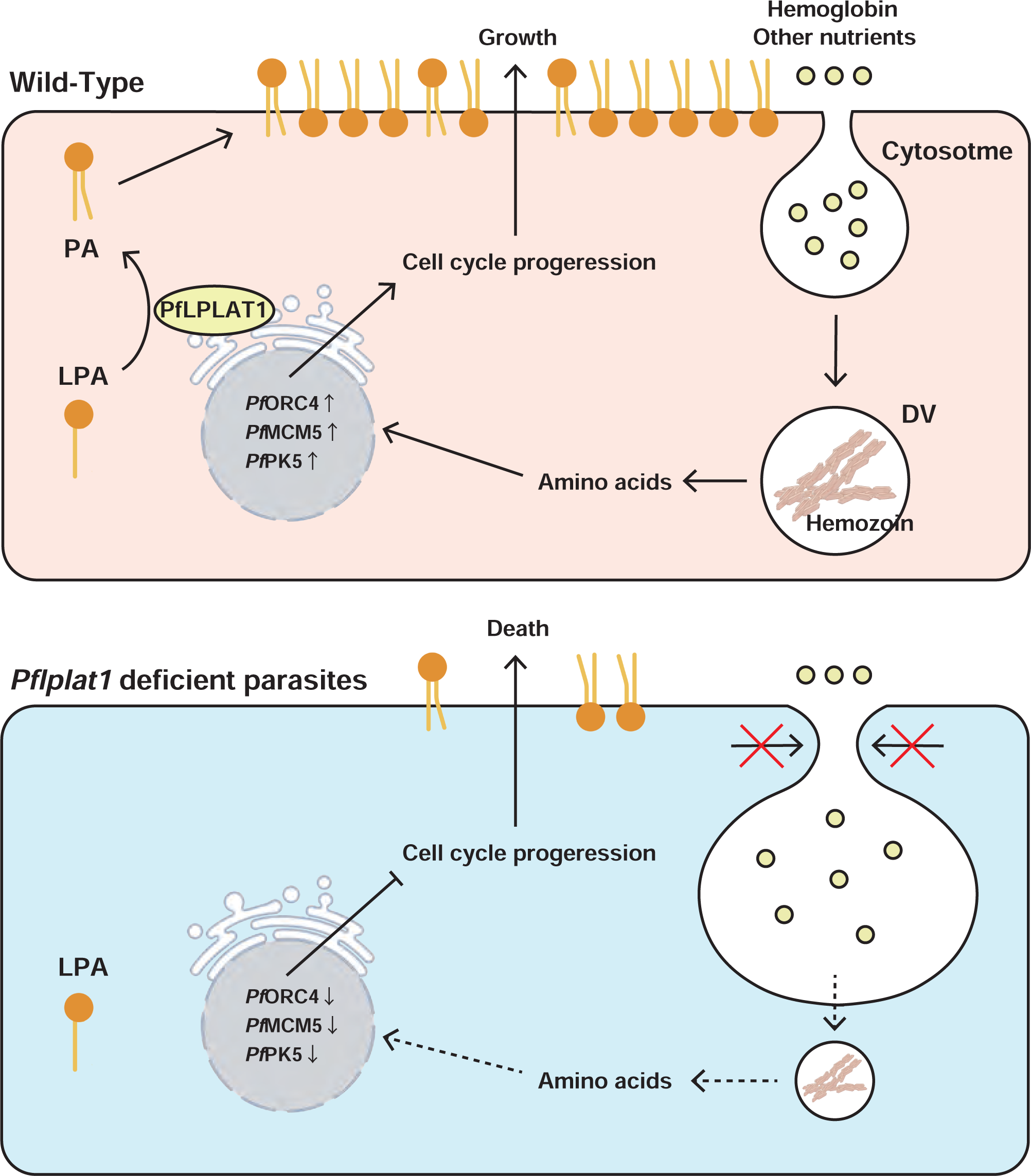
Schematic diagram showing the effect of *Pflplat1* deletion on parasites.

In mammals and yeasts, the cell cycle is rigorously regulated by mitotic kinases, including cyclin-dependent kinases (CDK), polo-like kinases, aurora kinases, NIMA-related kinases (NEK), protein phosphatases, and RING E3 ubiquitin ligases^54^. *P. falciparum* has some homologs of these proteins despite its tendency to become asynchronous with membrane-intact nuclear divisions followed by cytokinesis to separate multiple nuclei. Our transcriptomic analysis showed that PfCyc2, CDK-associated proteins, protein kinase 5 (PfPK5), and PfNEK3 were downregulated in *Pflplat1*-deficient parasites. Although the functions of most of these kinases remain unknown, PfPK5 was shown to be important for progression of parasites to the schizont stage through phosphorylation of origin recognition complex subunit 1 (PfORC1), a component of the pre-replication complex (pre-RC)^55^. Interestingly, *Pflplat1*-deficient parasites showed similar transcriptome profiles to those of dormant parasites, whose expression of pre-RC components (minichromosome maintenance complex component 5 (PfMCM5) and origin recognition complex subunit 4 (PfORC4)) and DNA mismatch repair proteins is decreased^56^. Therefore, *Pflplat1* deficiency may decrease the expression of pre-RC components and mitotic kinases, which retards the assembly of pre-RC and also prevents the disassembly of pre-RC. These phenomena might sequester parasites into the trophozoite stage and lead to the termination of cellular activities in parasites.

## Conclusion

Lipid metabolism and related areas in malaria parasites have been attractive for researchers as new study areas as well as potential drug targets^2^. Multi-membrane structures of infected erythrocytes and trafficking of both lipid and proteins are supported by lipid metabolism with a strict balance and molecular flux between organelles. Our study on PfLPLAT1 sheds light on the importance of the phospholipid synthesis pathway, particularly PA as an essential substance for the cell cycle progression of parasites. Considering our database search returned only a few entries that have sequence similarities, and PfLPLAT1 is an essential gene, other acyltransferases yet to be identified would function differently from PfLPLAT1 if they exist, but their roles should orchestrate each other to maintain normal cell cycle progression and parasitized erythrocyte structure.

## Methods

### Parasite culture

*Plasmodium falciparum* NF54::DiCre strain (kindly gifted from Moritz Treeck) was cultured in O^+^ erythrocytes at 5% hematocrit in RPMI-1640 media (Thermo Fisher Scientific, Waltham, MA, USA) supplemented with 5.95 mg/mL HEPES, 0.05 mg/mL hypoxanthine (Merck, Darmstadt, Germany), 2 mg/mL sodium bicarbonate (FUJIFILM Wako Chemical Corporation, Kanagawa, Japan), 10 μg/mL gentamicin (Thermo Fisher Scientific), and 0.5% Albumax II (Thermo Fisher Scientific) (RPMI-1640 complete media) and maintained at 37°C in a gas mixture of 5% O_2_, 5% CO_2_ and 90% N_2_, as previously described^7^. The experiments with human erythrocytes were conducted under the guidelines of the ethics committee of Nagasaki University (#191226226). Human erythrocytes were obtained from the Japan Red Cross Society (Tokyo, Japan) (No: R030027).

### Generation of *Pflplat1*:LoxPint:HA line

To conditionally knockout *Pflplat1*, we generated pSLI-*Pflplat1*-LoxP-3×2A-HA plasmid to introduce LoxP sites into *Pflplat1* utilizing a selection-linked integration (SLI) method^29^. To create pSLI-*Pflplat1*-LoxP-3×2A-HA, Pfef1a-5’UTR-pD3HA^57^ was linearized at BglII and XmaI restriction sites. Synthetic DNA containing a LoxP site followed by 3′ recodonized *Pflplat1*, triple influenza hemagglutinin (HA) sequences, a 2A self-cleaving peptide sequence, a neomycin resistance gene cassette, a LoxP site, and a GFP reporter gene was obtained (Azenta Life Sciences, Burlington, MA, USA). 5’ genomic *Pflplat1* was amplified from the genomic DNA of NF54::DiCre strain with CloneAmp HiFi PCR Premix (Takara Bio, Shiga, Japan) using primers listed in Table S1. These sequences were introduced into linearized Pfef1a-5’UTR-pD3HA using an In-Fusion HD Cloning Kit (Takara Bio).

For plasmid DNA transfection of pSLI-*Pflplat1*-LoxP-3×2A-HA, trophozoite and schizont-stage parasites were collected with MACS magnetic columns (Miltenyi Biotec, Bergisch Gladbach, Germany). RBCs were electroporated in cuvettes (Bio-Rad, Hercules, CA, USA) with the plasmid using a GenePulser Xcell (Bio-Rad). Collected parasites were incubated with electroporated RBCs in RPMI-1640 complete medium for 48 hours. After 48 hours, parasites were selected with 1 nM WR99210 (Merck) for 3–4 weeks followed by 400 µg/mL G418 (FUJIFILM Wako Pure Chemical Corporation) for 2 weeks. Limiting dilutions were performed using selected transgenic parasites to obtain clonal parasites. To confirm plasmid integration in the genome of clonal parasites, genomic DNA was extracted using a DNeasy Blood & Tissue Kit (QIAGEN, Venlo, Netherlands) and a diagnostic PCR was performed using the primers listed in Table S1.

### Generation of *Pflplat1*:LoxPint:HA^comp^

To generate the *Pflplat1* genetically complimentary line (*Pflplat1*:LoxPint:HA^comp^) from the *Pflplat1*:LoxPint:HA C9 line, we created a p*hsp86*-*Pflplat1*-myc rescue plasmid. To remove Cas9 coding sequence from pUF-Cas9-pre-sgRNA (Addgene plasmid #137845; http://n2t.net/addgene:137845; RRID:Addgene_137845), the plasmid was linearized at the XhoI and SalI restriction sites. Synthetic DNA containing *Pflplat1* coding sequence followed by sequences of linker and triple c-myc was obtained (Azenta Life Sciences). These sequences were introduced into linearized pUF-Cas9-pre-sgRNA using an In-Fusion HD Cloning Kit (Takara Bio).

For plasmid DNA transfection of p*hsp86*-*Pflplat1*-myc, trophozoite and schizont-stage *Pflplat1*:LoxPint:HA C9 were collected with MACS magnetic columns (Miltenyi Biotec). RBCs were electroporated in cuvettes (Bio-Rad) with the plasmid using a GenePulser Xcell (Bio-Rad). Collected parasites were incubated with electroporated RBCs in RPMI 1640 complete medium for 48 hours. After 48 hours, parasites were selected with 500 nM DSM-1 (Merck) for 3–4 weeks. To confirm that PfLPLAT1 was episomally expressed in *Pflplat1*:LoxPint:HA^comp^, we performed western blotting.

### Multiple sequence alignment of LPAATs

For multiple alignment of LPAAT sequences, the amino acid sequences were obtained from the NCBI database (https://www.ncbi.nlm.nih.gov/) (*Plasmodium falciparum* (*P. falciparum*) LPLAT1, XP_001348595.2; *Homo sapiens* (*H. sapiens*) LPLAT1, NP_001358367.1; *Mus musculus* (*M. musculus*) LPLAT1, AAB82009.1; *Caenorhabditis elegans* (*C. elegans*) ACL-1, CAB03160.1; *Saccharomyces cerevisiae* (*S. cerevisiae*) SLC1, NP_010231.1; *Toxoplasma gondii* (*T. gondii*), XP_002366705.1; *Babesia bovis* (*B. bovis*), XP_001610213.1; *Plasmodium yoelii* (*P. yoelii*) PY01678, XP_729473.2; *P. yoelii* PY02486, XP_730369.1; *Trypanosoma brucei* (*T. brucei*), XP_829573.1; *Escherichia coli* (*E. coli*), AAA24397.1). The alignment was performed using CLUSTALW (v2.1). Sequence logos were created using WebLogo (v2.8.2).

### Estimation of the three-dimensional structure of PfLPLAT1 and *Hs*AGPAT1

For three-dimensional construction of these enzymes, Protein Data Bank files were obtained from the AlphaFold Protein Structure Database (https://alphafold.ebi.ac.uk/) (PfLPLAT1; AF-Q8IL28-F1-model_v4, *Hs*AGPAT1; AF-Q99943-F1-model_v4). The three-dimensional structure was visualized using PyMoL (v0.99rc6).

### Growth assay

For growth assays, parasites were synchronized with 5% D-sorbitol (FUJIFILM Wako Pure Chemical Corporation) and seeded in 24-well plates. Synchronized parasites were cultured in 1 mL RPMI-1640 complete medium at 5% hematocrit and incubated with rapamycin (100 nM; Merck) or an equivalent volume of dimethyl sulfoxide. Parasitemia was counted every day using a Sysmex XN-30 hematology analyzer (Sysmex Corporation, Kobe, Japan)^58^ and kept lower than 5% by serial passages. The membranes of parasitized erythrocytes were permeabilized using CELLPACK DCL (Sysmex Corporation), and nucleic acids of parasites were stained using Fluorocell M (Sysmex Corporation) and Lysercell M (Sysmex Corporation). Growth curves were depicted using GraphPad Prism (v8.4.3) (GraphPad Software, Boston, MA, USA).

### Quantification of parasite stage distribution

For quantification of parasite stage distribution, trophozoite- and schizont-stage parasites were collected with MACS magnetic columns (Miltenyi Biotec). Collected parasites were incubated at 2% hematocrit in RPMI-1640 complete medium for 4 hours in agitated conditions (rotated at 40 rpm) to remove multiple infections of parasites in a single RBC. After 4 hours of incubation, parasites were tightly synchronized with 5% D-sorbitol (0-4 h post invasion) and seeded in six-well plates. Blood films were created at each time point and stained with Giemsa′s azur eosin methylene blue solution (Sigma–Aldrich, Burlington, MA, USA). Stage counting was performed manually under a light microscope.

### Immunofluorescence assay

Infected RBCs were immobilized on coverslips (Thorlabs, Inc., Newton, NJ, USA) coated with poly-D-lysine (MP Biomedicals, Santa Ana, CA, USA). After the immobilization, samples were fixed with 4% paraformaldehyde (Electron Microscopy Sciences, Hatfield, PA, USA) in phosphate buffered saline (PBS) for 10 min and 50 mM dimethyl suberimidate dihydrochloride (Sigma–Aldrich) in borate buffer for 20 min, followed by quenching with 10 mM glycine in PBS for 30 min. Fixed samples were permeabilized with 0.3% Triton X-100 (FUJIFILM Wako Pure Chemical Corporation) in PBS and blocked with 1% skim milk in 0.05% Tween 20 (Tokyo Chemical Industry Co., Ltd., Tokyo, Japan) in PBS for 60 min. Samples were treated with primary antibodies overnight and secondary antibodies for 60 min, followed by sealing with VECTASHIELD Antifade Mounting Medium with DAPI (Vector Laboratories, Inc., Newark, CA, USA). Mouse anti-HA antibody (6E2) (Cell Signaling Technology, Danvers, MA, USA) was used at 1:50 dilution. Rabbit anti-BiP antibody (MR4, Manassas, VA, USA) was used at 1:15 dilution. Alexa Fluor 488-conjugated goat anti-mouse (Thermo Fisher Scientific, Carlsbad, CA, USA), Alexa Fluor 594-conjugated goat anti-mouse, and Alexa Fluor 594-conjugated goat anti-rabbit antibodies were used at 1:1000 dilution. Samples were observed with a 100× objective using an LSM780 confocal laser scanning microscope (Carl Zeiss, Oberkochen, Germany).

ZEN lite (v3.5) (Carl Zeiss) was utilized to quantify the intensity of immunofluorescence signals in parasites. To plot signal profiles, the intensity on drawing lines was measured using the Profile Plot analysis tool in ImageJ (v1.53t) (https://imagej.nih.gov/ij/). For co-localization analysis, Pearson’s r values were calculated using the Coloc 2 tool in Fiji (v2.9.0) (https://imagej.net/software/fiji/).

### Western blotting

For western blot analysis, parasites were collected with MACS magnetic columns (Miltenyi Biotec) and lysed in 0.03% saponin solution for 5 min. Collected samples were dissolved in lysis buffer (4% SDS, 0.5% Triton X-100, 25-75 U DNase I (Takara Bio) and cOmplete protease inhibitor cocktail (Roche, Basel, Switzerland) in 0.5% PBS) and denatured with SDS sample buffer containing protein-reducing agents dithiothreitol. Lysates were boiled at 95°C for 3 min and run on polyacrylamide gels (Thermo Fisher Scientific). Proteins were transferred on PVDF membranes and blocked with a Bullet Blocking One (nacalai tesque, Kyoto, Japan). Blocked membranes were incubated with primary antibodies for 1 hour followed by secondary antibodies conjugated to horseradish peroxidase (HRP) (Jackson ImmunoResearch Laboratories Inc., West Grove, PA, USA) for 30 min. Mouse anti-HA antibody (6E2) (Cell Signaling Technology, Inc., Danvers, MA, USA) was used at 1:1000 dilution. Mouse anti-Myc antibody (9B11) (Cell Signaling Technology) was used at 1:1000 dilution. Mouse HRP was used at 1:5000 dilution. After probing with antibodies, the membranes were incubated with chemiluminescent HRP substrate (Thermo Fisher Scientific) for 5 min and proteins were visualized with WSE-6100 LuminoGraph I (ATTO, Tokyo, Japan). To investigate the protein expression levels of PfAldolase as an internal control, the membrane was washed with WB Stripping Solution Strong (nacalai tesque) for 30 min and blocked with Bullet Blocking One. After blocking, the membrane was incubated with HRP anti-plasmodium aldolase antibody (1:2000; Abcam, Cambridge, UK) for 30 min and chemiluminescent HRP substrate (Thermo Fisher Scientific) for 5 min.

### Gametocytogenesis

For gametocytogenesis, the parasite was maintained at 6% hematocrit in RPMI complete medium with 10% human serum (#28J0058; Japan Red Cross) and parasitemia was adjusted to 0.1%, as previously decribed^7^ (day 1). Hematocrit was reduced to 3% on day 4. The parasite was treated with 50 mM N-acetyl glucosamine (Sigma–Aldrich) to remove asexual parasites on days 9–11 and treated with 100 nM rapamycin (Sigma–Aldrich) from day 10. Giemsa-stained blood films were created to count gametocytemia and to investigate stage distribution of gametocytes manually on days 12–15. Media were replaced with pre-warmed and fresh medium every day up to day 15. For extraction of genomic DNA from the gametocyte, it was separated into an interface between layers of 40% and 60% Percoll and collected with MACS magnetic columns (Miltenyi Biotec) on day 14. Genomic DNA was extracted using a DNeasy Blood & Tissue Kit (Qiagen) and PCR was performed using the primers listed in Table S1.

### Transcriptomic analysis

For purification of RNA, parasites were collected with MACS magnetic columns (Miltenyi Biotec). Collected parasites were incubated at 2% hematocrit in RPMI-1640 complete medium for 4 hours in agitated conditions (rotated at 40 rpm) to remove multiple infections of parasites with a single RBC. After 4 hours of incubation, parasites were tightly synchronized with 5% D-sorbitol (0–4 h post invasion) and returned to RPMI-1640 complete medium with or without 100 nM rapamycin. Infected RBCs were harvested at 24 and 30 hours post-infection and dissolved in TRIzol Reagent (5 times the volume of RBCs) (Thermo Fisher Scientific). Dissolved samples were stored at −80°C until purification of RNA. RNA was separated from samples using Phasemaker Tubes (Thermo Fisher Scientific) and purified using a PureLink RNA Mini Kit (Thermo Fisher Scientific).

mRNAs were isolated from total RNAs using poly-T oligo-attached magnetic beads. For construction of a directional library, first-strand cDNA was synthesized using random hexamer primers and second-strand cDNAs were synthesized using dUTPs after fragmentation of captured mRNAs. After synthesis of double-stranded cDNA, end repair of double-stranded cDNA, 3’-adenylation (A-tailing), adapter ligation, size selection, USER enzyme excision, PCR amplification, and purification were performed. Real-time PCR and Qubit (Thermo Fisher Scientific) were utilized for quantification of the libraries. Quantified libraries were sequenced using Illumina Novaseq 6000 (Illumina Inc., San Diego, CA, USA). Raw data (raw reads in fastq format) were processed through in-house Perl scripts to remove reads containing adapters and ploy-N and reads with low quality. Paired-end reads were aligned to the reference genome (release-42 of *P. falciparum* 3D7) using Hisat2 (v2.0.5). Counting of the number of reads mapped to each gene was performed using FeatureCounts (v1.5.0). Differential expression analysis was performed using edgeR (v3.43.7). Adjusted p-values were calculated using the Benjamini–Hochberg method. Gene Ontology enrichment analysis of DEGs was performed using clusterProfiler (v4.9.1). The hypergeometric test was performed to calculate p-values.

### Hemozoin quantification

For quantification of hemozoin, parasites were collected with MACS magnetic columns (Miltenyi Biotec). Collected parasites were incubated at 2% hematocrit in RPMI 1640 complete medium for 4 hours in agitated conditions (rotated at 40 rpm) to remove multiple infections of parasites with a single RBC. After 4 hours of incubation, parasites were tightly synchronized with 5% D-sorbitol (0-4 h post invasion) and returned to RPMI-1640 complete medium with or without 100 nM rapamycin. Infected RBCs were observed under a Leica DMi8 inverted microscope (Leica Microsystems, Wetzlar, Germany) and exposed to polarized light to visualize hemozoins. The area of reflected lights (indicators of hemozoins) was measured using the Analyze Particle tool in ImageJ (v1.53t) (https://imagej.nih.gov/ij/).

### Sample preparation for transmission electron microscopy (TEM) and scanning transmission electron microscopy (STEM)

Sample preparation for TEM samples was slightly modified from previous papers ^59,60^. Briefly, the trophozoite and schizont stages of +Rap and −Rap parasites purified by MACS LS columns (Miltenyi Biotec.) were collected by centrifugation at 1300 × *g* for 5 min. These parasitized erythrocytes were fixed with 2% glutaraldehyde (Electron Microscopy Sciences) with 2% paraformaldehyde (Electron Microscopy Sciences) and 1% tannic acid (Merck) in 0.1 mol/L cacodylate buffer (TAAB Laboratories Equipment Ltd., Aldermaston, Berks, England), pH 7.4. Erythrocytes were further fixed with 1% osmium tetroxide aqueous solution (Electron Microscopy Sciences) for 1 hour at 4°C. Erythrocyte pellets were washed three times with ultrapure water, embedded in 2% agarose (PrimeGel Agrose LMT, Takara) when necessary, stained with 2% uranyl acetate (Electron Microscopy Sciences) on ice for 1 hour, and washed three times with ultrapure water.

The samples were serially dehydrated with 50%, 70%, 80%, 90%, 95%, and 100% ethanol for 10 min each. Infiltration of propylene oxide (FUJIFILM Wako Pure Chemical Corporation) was performed for 50 min at room temperature. Samples were embedded in propylene oxide with resin mixture; (23.8 mL Quetol-812, 11.2 mL DDSA, 15 mL MNA, and 0.9 mL DMP-30; Nisshin EM, Tokyo, Japan; 1:1 volume ratio) overnight. The samples were then heated for 30 hours at 60°C. Thick sections (approximately 250 nm) were generated with a diamond knife (DiATOME, Biel, Switzerland). After collecting sections by 200 mesh copper grids (GILDER GRID, No100; Gilder Grids Ltd., Grantham, Lincolnshire, UK), specimens were stained with 5% uranyl acetate solution (Merck) plus 0.4% lead citrate (TABB Laboratories Equipment Ltd), and then osmium coating was applied for observation by STEM (JEM-1000K RS, JEOL Ltd., Tokyo, Japan).

### Measurement of PfLPLAT1 enzymatic activities

To identify the specificities of both LPLs and acyl chain donors, 2 μg proteoliposomes was mixed with 25 μM of one type of deuterated LPLs (16:0-d9 LPA, 16:0-d31 LPC, or 16:0-d9 LPE) and a mixture of 1 μM fatty acid CoAs (16:0-CoA, 18:1-CoA, 18:2-CoA, 20:4-CoA, 20:5-CoA, and 22:6-CoA) in 100 mM Tris HCL (pH 7.4), 1 mM EDTA, and 2 mM CaCl_2_ (total reaction volume of 100 μL). The mixtures were incubated at 37°C for 30 min and the enzyme reaction was stopped by adding 500 μL MeOH. The samples were centrifuged, and the supernatants were transferred to new tubes.

The deuterated product diacyl phospholipids were detected by mass spectrometry and quantified based on the signal intensities of external standards for each lipid type (PC(34:1), PE(34:1), and PA(34:1)).

To prepare samples for lipid profile analyses, trophozoites and schizonts were separated by LS magnetic columns (Miltenyi Biotec), and further synchronized by incubating for 4 hours after mixing with 10 times the volume of non-parasitized erythrocytes and 5% sorbitol treatment. The number of cells were adjusted, and total lipids were directly extracted by treating synchronized parasites with 40 times the volume of MeOH.

### LC-MS analyses

Evaluation of enzyme activity and phospholipid profiles in the parasitized erythrocytes was performed by LC/ESI/MS/MS using a Nexera UHPLC system and a triple quadrupole mass spectrometer LCMS-8060 (Shimadzu Corp., Kyoto, Japan). An Acquity UPLC BEH C8 column (1.7 μm, 2.1 mm x 100 mm, Waters Corp., Milford, MA, USA) was used for chromatographic separation with three phases: 5 mM NH_4_HCO_3_/water (mobile phase A), acetonitrile (mobile phase B), and isopropanol (mobile phase C). Column oven temperature was set at 47°C. To analyze the enzymatic activity of PfLPLAT1 and PE and PC profiles in the parasitized erythrocytes, the pump gradient [time (%A/%B/%C)] was programmed as follows: 0 min (75/20/5)-20 min (20/75/5)-40 min (20/5/75)-45 min (5/5/90)-50 min (5/5/90)-55 min (75/20/5). For analyzing the PA profile in the parasitized erythrocytes, the pump gradient [time (%A/%B/%C)] was programmed as follows: 0 min (95/5/0)-8 min (70/30/0)-16 min (30/35/35)-28 min (6/47/47)-35 min (6/47/47)-35.1 min (95/5/0). Injection volume was 5 μL. Column oven temperature was set at 47°C and the flow rate was 0.35 mL/min. Multiple reaction monitoring transitions for phospholipids were set as follows: PC ([M+H] →184.1, positive mode), PE ([M+H] →[M+H]–141, positive mode), PA ([M-H] →153.1, negative mode), and SM ([M+H] →184.1, positive mode), where M is m/z of molecular-related ions. LabSolutions LCMS software (Shimadzu Corp.) was used for the operation of the equipment and data acquisition. Peak identifications and peak area integrations were performed using TRACES (https://github.com/KitaYoshihiro/TRACES)^61^. In profile analyses, area values of at least 10,000 were considered as reliable numbers, and lipid species with area values of more than 50,000 in the control groups were selected for the data analyses^61^.

### Statistical analysis

Data was checked for normality using the Shapiro–Wilk test. Statistical tests were performed using R software (v4.3.1) (https://www.r-project.org) and GraphPad Prism (v8.4.3).

## Data availability

The RNA sequencing data have been deposited in DDBJ Sequence Read Archive (DRA) (https://www.ddbj.nig.ac.jp/dra/index-e.html) as DRA017270.

## Acknowledgements

We thank Dr. Moritz Treek (The Francis Crick Institute, UK) for providing NF54::DiCre parasites for this study. We also thank Drs. Takao Shimizu (Institute of Microbial Chemistry, Tokyo, Japan), Eizo Takashima (Ehime University, Ehime, Japan), and Kazuhide Yahata (Ehime University, Ehime, Japan) for their helpful discussions and technical advice. We are also grateful for the technical assistance in high-voltage transmission electron microscopy and image analyses provided by Dr. Shigeo Arai, Ms. Tomoyo, M.S., and Ms. Nakao Kahoru Yoda, B.S., of the High-Voltage Electron Microscope Laboratory, Institute of Materials and Systems for Sustainability, Nagoya University, Japan. This work was supported partially by Grants-in-Aid for Scientific Research (KAKENHI) grant numbers: 17K08805 and 23K06518 (F.T.). The Department of Lipidomics, Graduate School of Medicine, University of Tokyo, is supported by Shimadzu Corp. (Kyoto, Japan). The Department of Cellular Architecture Studies, Institute of Tropical Medicine, Nagasaki University, is supported by Shionogi and Co., Ltd. These funders were not involved in the design, experiments, or data analyses for this study. Finally, we thank H. Nikki March, PhD, from Edanz (https://jp.edanz.com/ac) for editing a draft of this manuscript.

## Author contributions

Conceptualization: J.F., F.T., S.M., and H.S. Data collection: J.F., S.M.T., M.Y., E.S.H., J.U., and F.T. Data analysis: J.F., S.M.T., M.Y., E.S.H., T.S., D.K.I, J.U., H.S., and F.T. Funding acquisition: F.T. Supervision: K.K., J.U., H.S, and F.T. Writing – original draft: J.F. and F.T. Writing – review & editing: J.F., S.M.T., M.Y., E.S.H., S.M., T.S., D.K.I., K.K., J.U., H.S., and F.T. Resources: F.T. and K.K.

## Supplementary materials

**Figure S1. Plasmid DNA integration and *Pf*lplat1-knockout efficiency in *Pflplat1*:LoxPint:HA G6 clonal line**

(A) Diagnostic PCR was performed to check whether the construct was correctly integrated into the target locus. DNA segments were amplified with f1/r1 (NF54::DiCre, not amplified; *Pflplat1*:LoxPint:HA G6, 2676 bp), f2/r2 (NF54::DiCre, not amplified; *Pflplat1*:LoxPint:HA G6, 979 bp), and f1/r2 (NF54::DiCre, 1027 bp; *Pflplat1*:LoxPint:HA G6, not amplified) primer sets. (B–D) Conditional knockout of *Pflplat1* was confirmed by PCR, western blotting, and IFA. (B) The size of DNA amplicons was shifted from 2676 to 784 bp at 24 hours of rapamycin treatment. (C) A reduction in PfLPLAT1 expression was detected in *Pflplat1*:LoxPint:HA G6 with anti-HA antibody at 24 and 48 hours of rapamycin treatment. PfAldolase was used as an internal control. Experiments were repeated three times. (D) Left panel: representative immunofluorescence micrographs of *Pflplat1*:LoxPint:HA G6 at 24 hours of rapamycin treatment. Samples were stained for nuclei with DAPI (cyan) and PfLPLAT1 with anti-HA antibody (magenta) and scanned using confocal microscopy. Scale bars = 2 μm. Right panel: violin plots showing the fluorescence intensity of HA-tagged PfLPLAT1 in rapamycin-treated (+Rap) or untreated (−Rap) *Pflplat1*:LoxPint:HA G6 (n = 20 per group). Violin plots range from minimum to maximum values. Dotted bottom, middle, and top horizontal lines show 25th percentile, median, and 75th percentile. Experiments were repeated two times. Statistical analysis was performed using F-test for equality of variance followed by two-tailed t-test.

**Figure S2. *Pflplat1* deletion did not influence gametocyte maturation**

(A) Conditional knockout of *Pflplat1* in gametocytes was checked by performing PCR at day 14 after gametocytogenesis. The size of DNA amplicons was shifted from 2676 to 784 bp. (B) Gametocytemia of +Rap or −Rap *Pflplat1*:LoxPint:HA on days 12–15 after gametocytogenesis. Data are shown as mean ± SD from n = 3 independent biological replicates. (C) Heatmap showing gametocyte stage distribution of +Rap and −Rap *Pflplat1*:LoxPint:HA. Data are shown as mean from n = 3 independent biological replicates.

**Figure S3. *Pflplat1* deletion reduces the size of hemozoin in parasites**

Left panel: violin plots showing the size of hemozoin in +Rap or −Rap *Pflplat1*:LoxPint:HA (n = 60 per group). Violin plots range from minimum to maximum values. Dotted bottom, middle, and top horizontal lines show 25th percentile, median, and 75th percentile. Data was combined from three independent experiments (n = 20 per group per experiment). Statistical analysis was performed using the Wilcoxon signed-rank test. Right panel: representative polarized light micrographs of *Pflplat1*:LoxPint:HA at 30 hpi. Images were obtained with “crossed polars” and a 100× HCX PL FLUOTAR objective (n = 1.32)-equipped Leica DMi8 fluorescence microscope. Scale bars = 2 μm.

**Figure S4. Transcriptomic analysis of transgenic parasites at 24 and 30 hpi**

(A) Volcano plot showing DEGs in +Rap versus −Rap *Pflplat1*:LoxPint:HA at 24 hpi (n = 3 independent biological replicates per group). Blue dots indicate downregulated genes (log_2_FC < 0.5, adjusted p-value < 0.05). Dotted vertical lines show 0.5 and −0.5 of log_2_FC and dotted horizontal line shows 0.05 of the adjusted p-value. Adjusted p-values were calculated using the Benjamini–Hochberg method. (B) Principal component analysis of gene expression profiles in +Rap and −Rap samples at 24 and 30 hpi. (C) Heatmaps showing the Pearson correlation coefficient between gene expression profiles in +Rap and −Rap *Pflplat1*:LoxPint:HA at 24 and 30 hpi.

**Figure S5. *Pflplat1* deficiency affects the profile of phospholipids in parasites**

(A–D) Bar graphs showing normalized fractions of PA (A), PE (B), PC (C), and SM (D) against the total area values within the same type of lipid in +Rap and −Rap *Pflplat1*:LoxPint:HA at 30 hpi. Data are shown as mean ± SD from n = 3 independent biological replicates. P-values were calculated using unpaired two-tailed t-test (*, *P* < 0.05; **, *P* < 0.01, ***, *P* < 0.005).

**Table S1. Primers used in this study**

**Table S2. DEG list and GO analysis results**

