## Supplementary figures and images for "Pivotal roles of *Plasmodium falciparum* lysophospholipid acyltransferase 1 in cell cycle progression and cytostome internalization"

### Supplementary figures

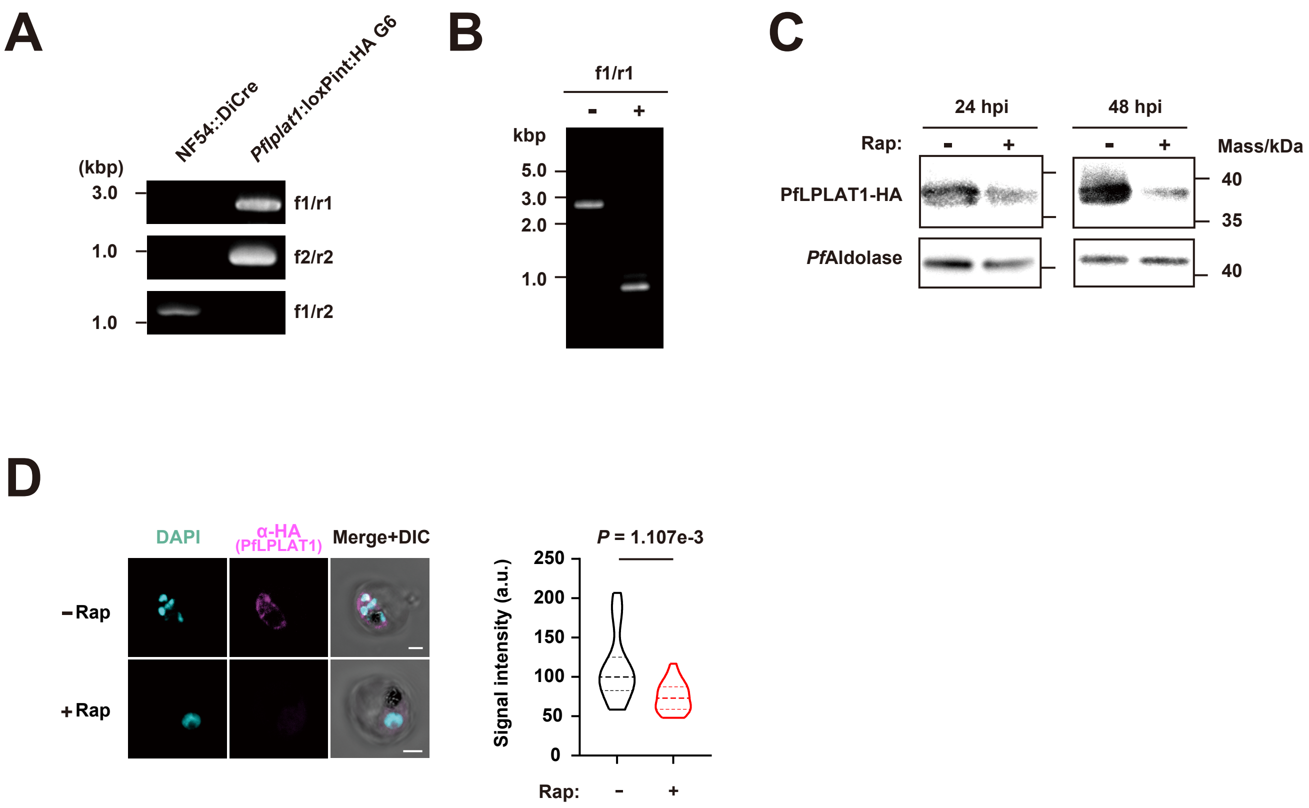

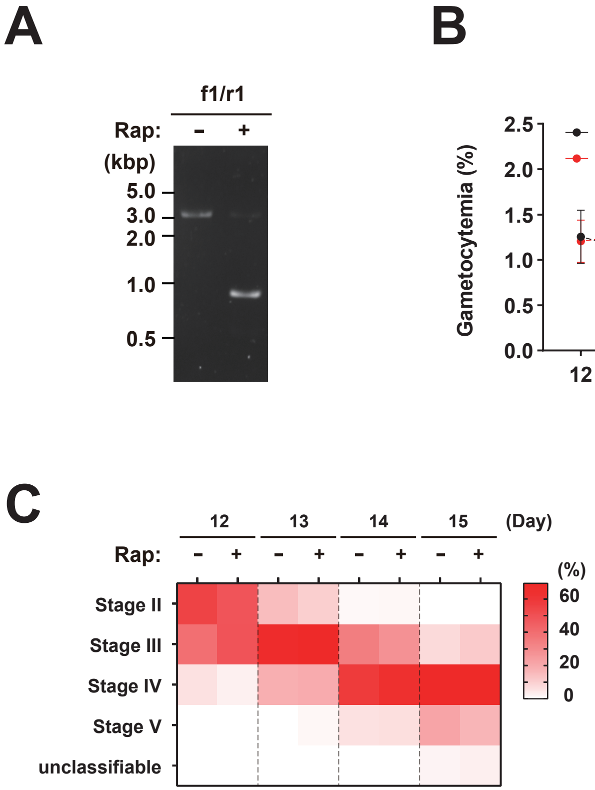

#### Figure S3

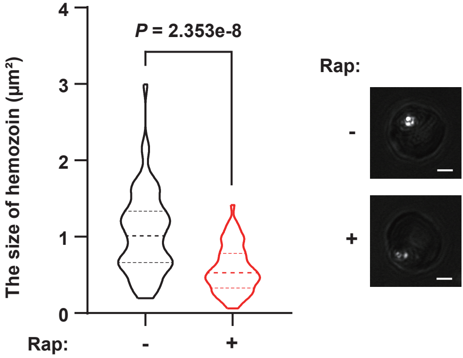

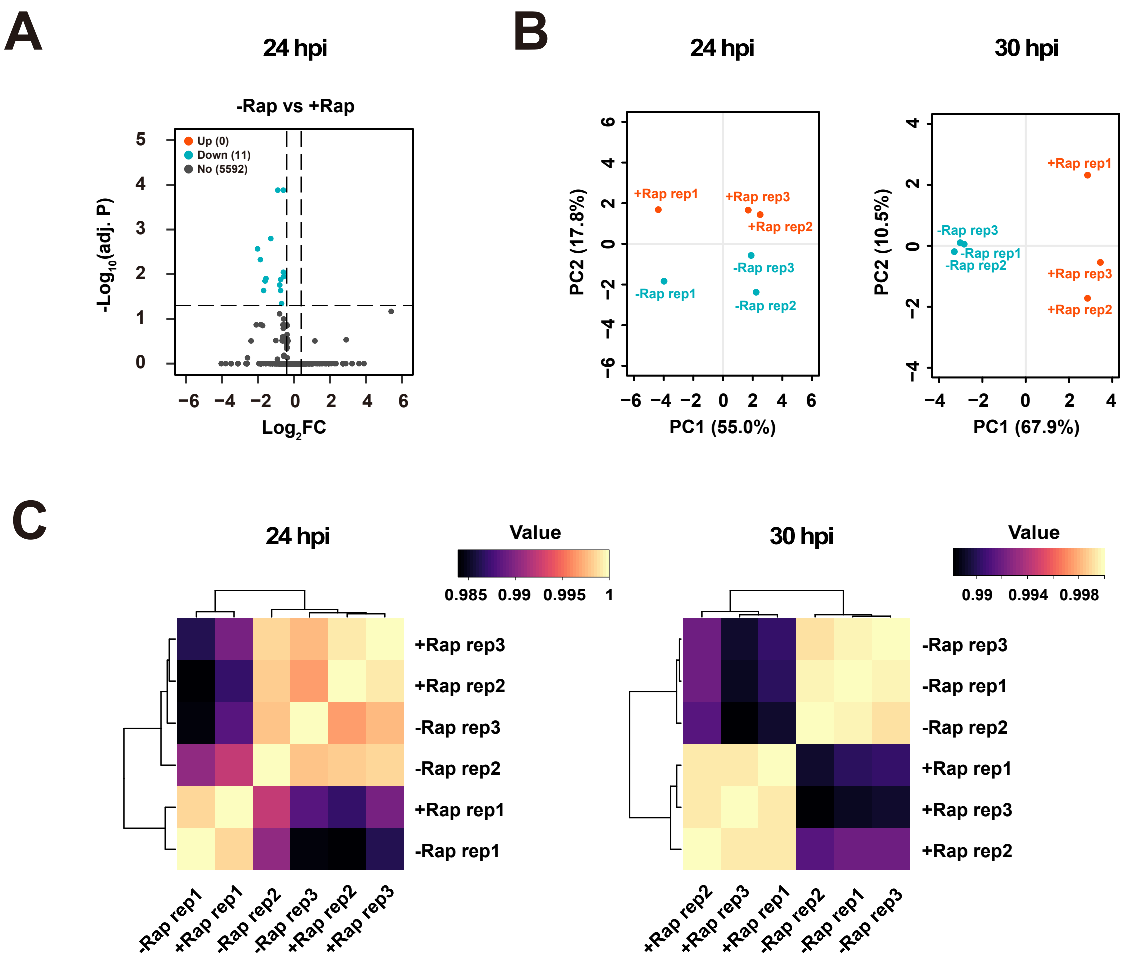

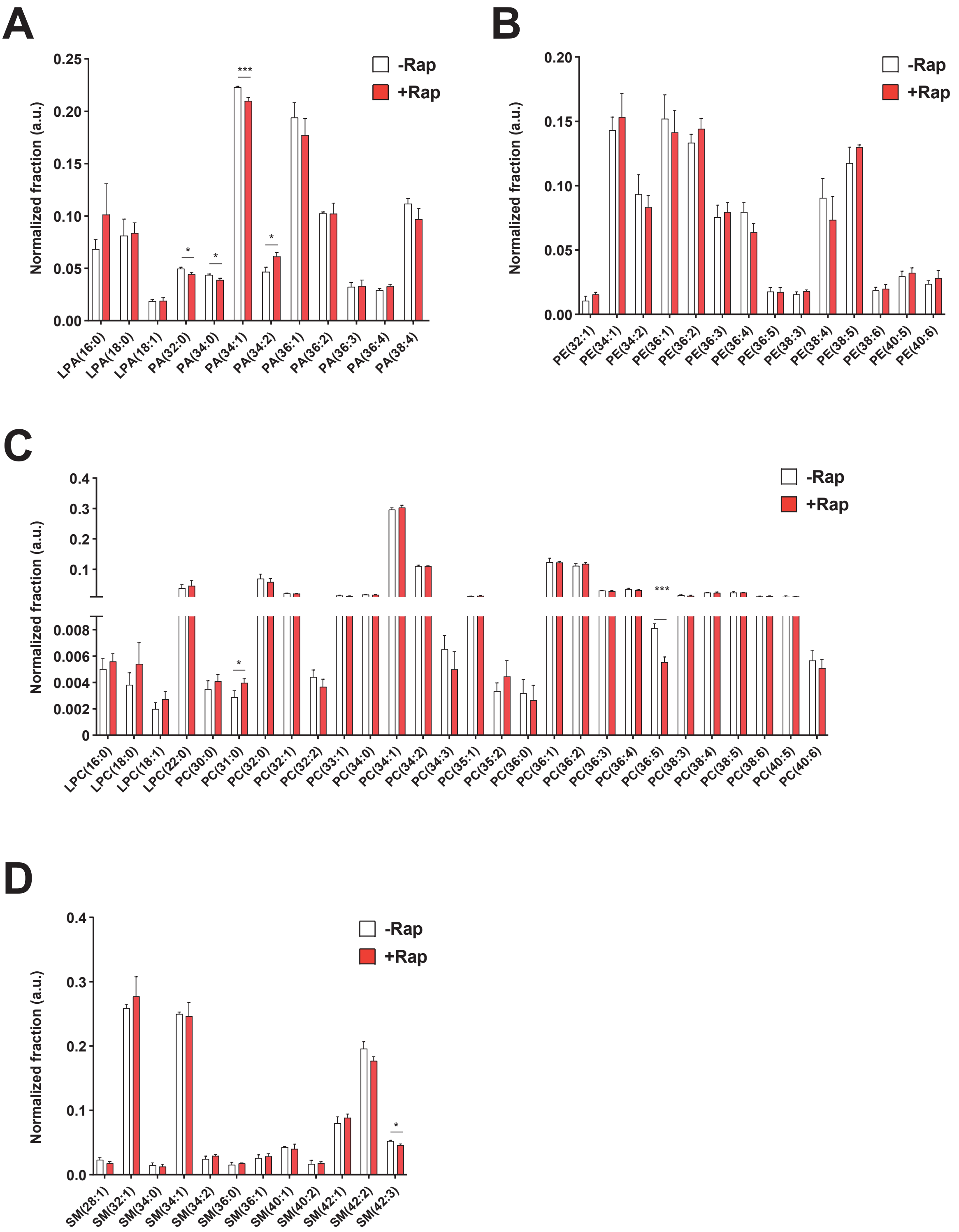
